# Age-trajectory of mother-infant relationships in wild Assamese macaques

**DOI:** 10.1101/2025.05.10.653242

**Authors:** Ana Lucia Arbaiza-Bayona, Roger Mundry, Suchinda Malaivijitnond, Suthirote Meesawat, Oliver Schülke, Julia Ostner

**Affiliations:** Department of Behavioral Ecology, Georg-August-Universität Göttingen, Johann-Friedrich-Blumenbach Institute for Zoology and Anthropology, Kellnerweg 6, D-37077 Göttingen, Germany; Research Group Primate Social Evolution, German Primate Center, Leibniz Institute for Primate Research, Kellnerweg 4, 37077 Göttingen, Germany.; Leibniz ScienceCampus Primate Cognition, German Primate Center, Leibniz Institute for Primate Research, Kellnerweg 4, 37077 Göttingen, Germany.; Department for Primate Cognition, Georg-August-Universität Göttingen, Johann-Friedrich-Blumenbach Institute, Kellnerweg 4, 37077 Göttingen, Germany.; Cognitive Ethology Laboratory, German Primate Center, Leibniz Institute for Primate Research, Kellnerweg 4, 37077 Göttingen, Germany.; National Primate Research Center of Thailand, Chulalongkorn University, Saraburi, Thailand.; Department of Biology, Faculty of Science, Chulalongkorn University, Bangkok, Thailand.

**Keywords:** mother-infant spatial relationship, infant independence, maternal investment, primates, development

## Abstract

Maternal care is ubiquitous in mammals, yet its degree and duration vary across taxa. Primate mothers provide extended care, with similar developmental transitions of the mother-infant relationship, though with different paces of change. Ecological conditions can influence the trajectory of this relationship, but data from the wild are still scarce. We used methods from growth studies to quantitatively describe the non-linear age-trajectory of the mother-infant spatial relationship, and the transition from dependent to independent feeding and locomotion in wild Assamese macaques (*M. assamensis*). We also explored sex differences in the development of the mother-infant relationship. We used a modified Gompertz function to model the combined effect of infant age and sex on mother and infant behaviors extracted from focal observations of 58 infants. Newborns were fully dependent on their mothers for feeding and transportation, with mothers maintaining close proximity. A transitional phase emerged between 1 and 3 months of infant age, marked by a noticeable reduction in the spatial proximity with the mother and a shift in the responsibility for the infant’s feeding and transportation. During the second half of infancy, the decrease in proximity time slowed down, with infants achieving near-complete locomotion independence, spending the majority of time away from their mothers and feeding independently. No sex differences were found. Our models provided a robust fit for most variables, but we recommend future exploration of alternative nonlinear functions. We interpret the early infant independence observed in our population in the context of the species reproductive strategy.

## Introduction

Parental care is widespread in vertebrates but varies considerably among taxa in terms of who provides it, as well as how and for how long it is given (Rosenblatt, 1998). Mammalian females invest heavily in gestation and lactation, with varying degrees and forms of postnatal care. Primates, in particular, have a slow life history for their body size compared to other mammals (Charnov & Berrigan, 2005) and are characterized by prolonged maternal care. Primate mothers may extend their care even beyond weaning, sometimes continuing into later life stages (van Noordwijk, 2012), offering ample opportunities for maternal influence on offspring behavior (Maestripieri & Mateo, 2009). The interaction between an infant primate and its mother begins with the physiological connection established from conception and develops after birth into a close relationship that provides nutrition, protection, thermoregulation, and means of transportation (Fairbanks, 2003; Nicolson, 1991). Moreover, the mother serves as a source of learning by offering safe opportunities for direct social learning (Mikeliban et al., 2021; Sargeant & Mann, 2009), and by acting as a secure base from which infants can explore and learn about their ecological and social environment (Ainsworth, 1979; Suomi, 2005; Uomini et al., 2020)). Thus, primate mothers provide intensive and extended care to their infants, who rely on them for survival until they develop the necessary skills to become self-sufficient.

When a primate is born, its mother maintains close contact and proximity, dedicating considerable time to lactation and carrying (Altmann & Samuels, 1992; Rosenblatt, 1998). This significant investment requires adjustments of the mother’s activity budget to accommodate the corresponding increase in energy expenditure (Dittus & Baker, 2024; Dunbar & Dunbar, 1988; Touitou et al., 2021). As the infant matures and develops the necessary motor skills to move away from its mother, it begins to break contact and explore the ecological and social surroundings (Byrne & Suomi, 1995). However, to mitigate the risk of mortality from predation, intragroup aggression, or injury, the mother initially restrains her infant’s attempts to move away and actively minimizes the distance sought by the offspring (R. A. Hinde & Spencer-Booth, 1968; Lycett et al., 1998; Maestripieri, 1995). Over time, the mother decreases the energetic investment in the infant, shifting energy allocation more towards her own maintenance and future reproduction (Trivers, 1974). The mother gradually increases the distance between herself and her infant, while refusing the infant’s attempts to nurse or to be carried for transportation. In response, the infant tries to maximize its own growth and survival by displaying signs of distress to regain maternal care or by actively seeking contact and proximity with her (Maestripieri, 1995). The transition from infant dependence to independence, therefore, involves a delicate balance: the mother protects the infant to ensure its survival while simultaneously promoting its independence to optimize her future reproductive success and survival.

The changes of the mother-infant relationship described above are shared among different primate taxa, though the pace of moving through these changes varies significantly between species and populations (Harvey & Clutton-Brock, 1985; Langer, 2003; Young & Shapiro, 2018). For instance, the timing of locomotory independence, defined as infants being carried less than 20% of their daily traveling time, can vary from 35 days after birth in Senegal bushbabys (*Galago senegalensis*) to 1,826 days after birth in chimpanzees (*Pan troglodytes*) and bonobos (*Pan paniscus*) (Young & Shapiro, 2018). Life-history theory provides a framework to understand this variation (Stearns, 1976), whereby smaller species and those with smaller relative brain size generally develop faster, and larger species and those with larger brain size develop more slowly (Barrickman et al., 2008; Lee et al., 1991; Sol, 2009). Yet, life history traits are not only explained by allometric features but also by ecological factors such as the extrinsic mortality level (Promislow & Harvey, 1990) or the diet of a given population (Borries et al., 2011; Janson & van Schaik, 1993; Robbins et al., 2023). For example, Asian colobines and macaques in the wild exhibit longer gestation periods, longer interbirth intervals, and a later age at first birth compared to provisioned populations of the same species (Borries et al., 2011). Likewise, differences in the developmental speed of lowland and mountain gorillas (*G. gorilla* and *G. beringei*) have been explained by variations in their feeding ecology (Robbins et al., 2023). That ecological conditions affect life-history traits, highlights the importance of studying different populations exposed to diverse ecological demands. The genus *Macaca* has been the focus of pioneering work on mother-infant relationships (R. A. Hinde and Davies, 1972; Maestripieri, 1995; Thierry, 1985), but behavioral data from primate populations living in their natural habitats are still scarce. Here, we provide data on Assamese macaques (*M. assamensis*) living in their natural habitat and use methods from growth studies to quantitatively describe temporal changes in their mother-infant relationship.

Assamese macaques are female philopatric and conceive their first offspring at around 5.5 years of age (Anzàa et al., 2022; Fürtbauer et al., 2010). They live in multimale-multifemale groups with a promiscuous mating system (Fürtbauer et al., 2010; Ostner et al., 2008) and reproduce seasonally with interbirth intervals of either 1 or 2 years (Fürtbauer et al., 2010; Shivani et al., 2025). Our aim was to explore the age dependent trajectories of the mother-infant relationship in the population in terms of (a) the spatial behavior of mothers and their infants towards each other and (b) the transition from dependent to independent feeding and locomotion. Given the altricial nature of primates, we expected high amounts of maternal care and minimal infant independence immediately after birth, when newborns are highly dependent on their mothers for feeding and transportation, and spend the majority of their time in body contact or close proximity to their mother. This would be followed by a gradual decline in mother proximity and body contact seeking behaviors, accompanied by a steeper decrease in high-investment maternal behaviors (i.e., carrying and nursing). At the same time, we expected a corresponding increase in independent locomotion and feeding by the offspring. While we hypothesized that high-investment maternal behaviors would cease by the end of infancy, driven by an increasing rate of maternal rejection of infant attempts to initiate contact, we anticipated some level of mother-infant proximity and body contact to stabilize during the later stages of infancy. Thus, because we did not expect a linear age-trajectory in maternal and infant behaviors, we used non linear modelling techniques to quantitatively describe the developmental transitions in the mother-infant relationship.

We further investigated whether infant sex influenced the developmental trajectory of mother-infant relationships in our study population. In philopatric species, mothers tend to invest more in the resident sex due to the increased potential benefits of maintaining a social bond with individuals who will remain in the group. For example, in female-philopatric rhesus macaques, mothers are more aggressive and less likely to groom or maintain body contact with male compared to female infants (Kulik et al., 2016). In contrast, in species characterized by male philopatry, such as black handed spider monkeys (*Ateles geoffroyi*, Symington, 1987) and chimpanzees (Băadescu et al., 2022), mothers tend to invest more in their male compared to their female offspring. Given the mandatory high maternal investment required during the initial ontogenetic phase to ensure infant survival, we expected sex differences in maternal care and infant independence to become more pronounced with infant age. Based on the dispersal regime of the studied species, we expected mothers of male infants to decrease their carrying, nursing, and contact and proximity-seeking behaviors earlier and/or faster than mothers of female infants. In conjunction, we expected that male infants exhibit an earlier increase in body contact and proximity-seeking behaviors as well as their feeding and locomotion independence, compared to female infants.

## Methods

### Study site and subjects

The study was carried out on a population of Assamese macaques at the study site XXX, that has been followed since 2006. The study area consists of hill evergreen forest with patches of bamboo forest and large populations of large herbivores and predators as an indication of its low level of disturbance (Borries et al., 2011). Reproduction is seasonal with ∼ 80 of births occurring in April-June and most conceptions between November-January (Fürtbauer et al., 2010; Shivani et al., 2025; Touitou et al., 2021). Despite seasonal reproduction, environmental predictability is low with high variation in fruit availability within and between years (Berghäanel et al., 2016; Heesen et al., 2013). Infancy was defined as the first 12 months of life, given that females can give birth to a new offspring the year after the last offspring was born (Shivani et al., 2025) and nipple contact is observed during the first year of age, despite a significant reduction around 6 months of infant age (Berghäanel et al., 2016).

In this study, we focused on 58 infants born in three cohorts (2011, 2018 and 2022) to 40 different mothers, with a sex distribution of 34 male (59%) and 24 female (41%) infants. Of the 40 mothers, 10 were primiparous (25%), and 15 mothers had offspring in more than one cohort (12 in two cohorts, three in three). Individuals were identified based on physical features, and relatedness of mother-offspring dyads was determined through observations of nipple contact during the infant’s first weeks of life. Of the 58 infants, all but four female infants survived until the end of their first year of life. These female infants were included in the analysis since they disappeared late (at 5 and 7 months of age), but one male infant that was 1 month old when disappeared was excluded from the analysis. We had nine combinations of group and year in our data, which resulted from various splits of the original groups across years. In 2011 one study group was followed, in 2018 three and in 2022 five. Between-group variation in size ranged from 18 to 74 individuals, but within group variation per year was relatively low (*m*=8.875, *sd* =2.9).

### Data collection

We observed infants from birth onwards using 40-min continuous and instantaneous focal sampling (Altmann, 1974) to record mother-infant interactions. For the instantaneous sampling, we collected sampling points with a 2-minute interval. In 2018 and 2022, we spent approximately one block of 5 to 10 days per month with each group. Thus, for the cohorts of 2018 and 2022, we used the block as the temporal unit for data aggregation. For the 2011 cohort, we used the calendar month as the temporal unit. This resulted in 3 to 13 data points (combinations of infant and block) for each mother-infant dyad (*m*=8.82, *sd* =2.24), depending on the survival of the infant. Our total observation time was 2,255 h, with a mean observation time of 4.40 h (*sd* =2.35 h) per individual and block. This represented 25,283 sampling points of the instantaneous sampling, with a mean number of sampling points of 150 (*sd* =83.2) per individual and block. Within each data collection period, we conducted focal animal observations by two or more observers and compared and discussed parallel protocols to promote inter-observer reliability.

Based on the long-term project’s ethogram and considering the variables commonly used to describe mother-infant relationships in primates (e.g., Arbaiza-Bayona et al., 2022; Bardi and Huffman, 2002; Deng and Zhao, 1991; Revathe et al., 2024), we recorded the time and direction of initiation and termination of body contact and proximity between infants and mothers, as well as events in which the mother restrained her infant’s movement or refused the infant’s attempts to establish body contact, all in the continuous sampling protocols. We were able to record the initiation and termination of mother-infant body contact and proximity in 55% and 73% of all cases, respectively. In the instantaneous part of the protocol, we recorded whether the infant was in nipple contact with the mother, being carried by the mother, or moving or feeding independently (see Table S1 for the definition of each behavior).

### Model formulation

#### Infant age

The age of the infant was included in the model as the main predictor of maternal and infant behavior. The day of birth of the infant was either known to the exact day or was calculated as the mid-point between the last day the mother was seen without an infant and the first day the infant was seen. Exact date of birth was known for 22 of the 58 infants; for the remaining infants the median error around the date of birth was 14.25 days (*Q1* =2 days, *Q3* =16 days, range = 2:24). Given that we aggregated the data per block, we used the mean age of the infant on months during the block.

Based on our expectations regarding the age-related trajectories of the mother-infant spatial relationship and the infant’s transition from dependent to independent feeding and locomotion, we predicted a nonlinear relationship between age and both infant and maternal behaviors, which we modeled using a Gompertz function (Gompertz, 1833). The Gompertz function is a sigmoid function, which unlike the logistic function, allows for an asymmetric approach to the curve’s initial and final asymptotes. In addition, other than a standard logistic model, it allows for a lower asymptote larger than 0 and an upper asymptote smaller than 1. Although the Gompertz function was originally developed to explain changes in mortality rates as a function of human age (Gompertz, 1833), it has been applied to model temporal changes in other biological processes, such as body growth in mammals (Zullinger et al., 1984). Also, the Gompertz function has been used to fit primate data on the ontogenetic trajectory of time spent in suckling and out of contact with caretakers (Tab Rasmussen & Tan, 1992).

We modified the original function by adding a constant representing the initial or left asymptote (*d*):

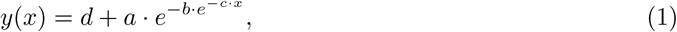

where:

*y* is the variable representing maternal or infant behavior,

*d* is the initial or left asymptote,

a. is the amplitude of change, determining the difference between *d* and the final or right asymptote of *y*(*x*),
b. determines the displacement along the time axis, together with *c* controlling at which value of *x* the steepest change of *y* occurs,
c. is the rate parameter that controls how quickly *y*(*x*) transitions between its asymptotes, and *x* represents infant age, the independent variable driving changes in *y*(*x*).

This function models the increase in infant independence (*y*) with respect to infant age (*x*) (i.e., *a >*0), whereas a decrease with age was expected for maternal care (i.e., *a <*0), and hence instead of adding (*a*) to (*d*), we subtracted (*a*) from (*d*).

#### Fruit availability

Since Assamese macaques reproduce seasonally, we wanted to control for environmental conditions that vary with offspring age and could also influence infant and maternal behavior. Specifically, given that Assamese macaques in this population are predominantly frugivorous (Schülke et al., 2011), we included an index of population level fruit availability as a potential confounding variable in our models, which has been shown to correlate with female energy intake, behavioral changes, and hormonal levels (Heesen et al., 2013). Fruit abundance was measured monthly in several individuals of 30 tree species that in more than one month accounted for more than 5% of the population’s monthly fruit feeding time and collectively attracted more than 72% of the population’s total fruit feeding time over a 10-year study period. For each tree species *i*, a monthly index was calculated by combining: (a) the average abundance of fruits across the monitored individuals of the species (*A_i_*), (b) the estimated density of the species over 20.25 ha of botanical plots within the study area (*D_i_*), and (c) the average basal area of the species (*B_i_*). The resulting values for the 30 species (*n*) were summed to calculate the monthly Fruit Availability Index (FAI) as Equation 2.

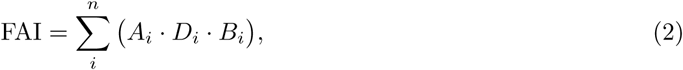

Daily FAI scores were then interpolated from the monthly values, allowing us to assign an average daily FAI to each focal block.

#### Maternal characteristics

Maternal parity and dominance rank can lead to variations in maternal care among females (Altmann & Samuels, 1992; Berman et al., 1994; Liu et al., 2024; Roura-Torres et al., 2025; Tanaka, 1989). Although our goal was not to explain the potential effects of these variables on the mother-infant spatial relationship and the infant’s transition from dependent to independent feeding and locomotion, we included them as additional fixed effects to account for biases arising from our unbalanced sampling (e.g., we could not choose which mothers gave birth in the studied cohorts) and to improve the precision of the models. We assumed that there is no true causal relationship between parity and maternal rank and any of the other variables included in the model. Given that previous studies suggested an effect of maternal rank and parity on maternal and not infant behavior, the models for independent feeding and locomotion did not include maternal parity and rank. However, because the spatial relationship between the mother and infant is influenced by the behavior of both members of the dyad, both variables were included in the models analyzing the infant-initiation of body contact and proximity with their mothers.

To estimate the dominance rank of the 40 mothers we used the Elo-rating method (Albers & De Vries, 2001) using the elo.seq function from the EloRating R package (Neumann & Kulik, 2020) with 1 000 as the starting value and 100 as the gain constant *k* (De Moor et al., 2019; Stranks et al., 2024). Data points were decided dyadic conflicts between females during which the losing female exhibited only submissive behaviors (bared-teeth display, make-room, give-ground), while the winner showed no submissive behaviors. The winner either displayed only aggressive behaviors (open-mouth threat, lunge, slap, bite, chase) or did not actively provoke the submission. The Elo-rating calculations for each group included the history of wins and losses from all recorded conflicts. Thus, although Elo-scores were calculated based on the individuals present in each group at a given moment, they were still influenced by the history of agonistic events prior to the group split. We used standardized Elo-scores (assigning 1 to the highest and 0 to the lowest score per group, see Table S2) at the beginning of each birth season to represent the relative rank of each mother within the group.

### Data analysis

We excluded data points for which we had less than 1 hour of observation time for an infant in a given block to aim for representative data on infant and mother behavior. This left 512 data points in total. However, models using instantaneous sampling data included 511 data points because for one combination of block and infant we had no data. Also, for some models in which the response variable was a quotient, data points were excluded when the total number of occurrences in the denominator was 0 (e.g., no proximity changes), reducing the total number of data points used to fit the model (see Table 1).

**Table 1.**
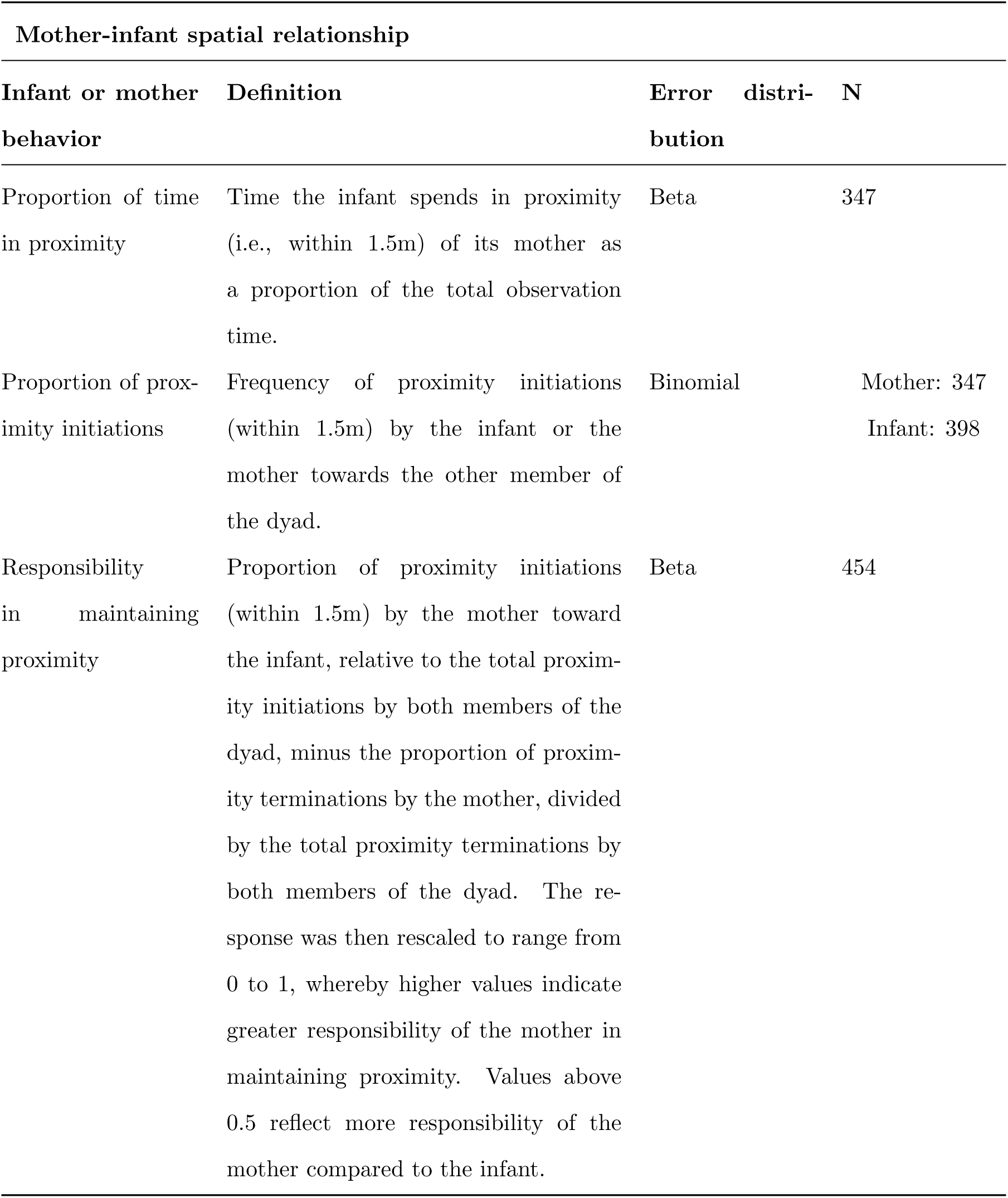
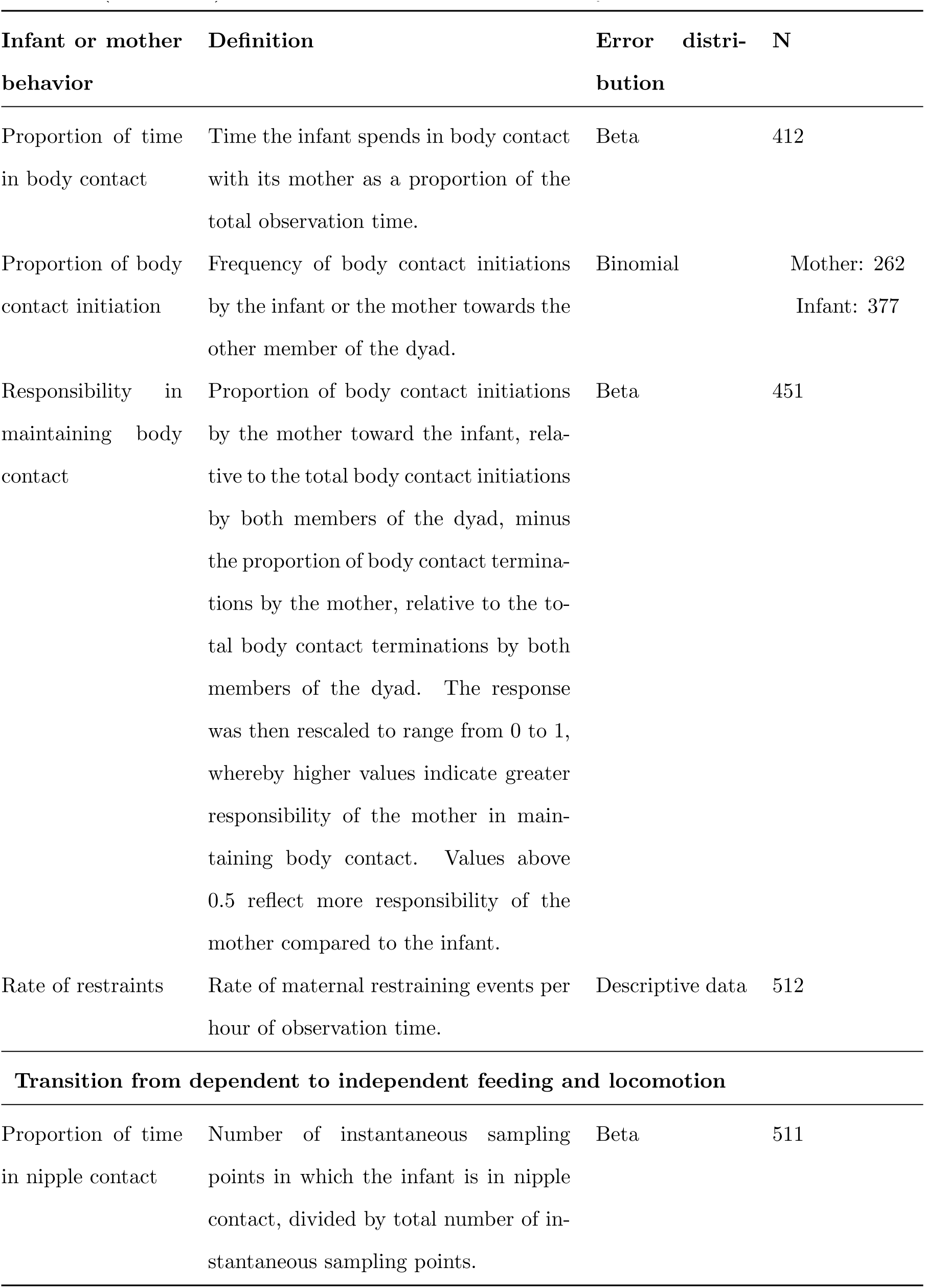
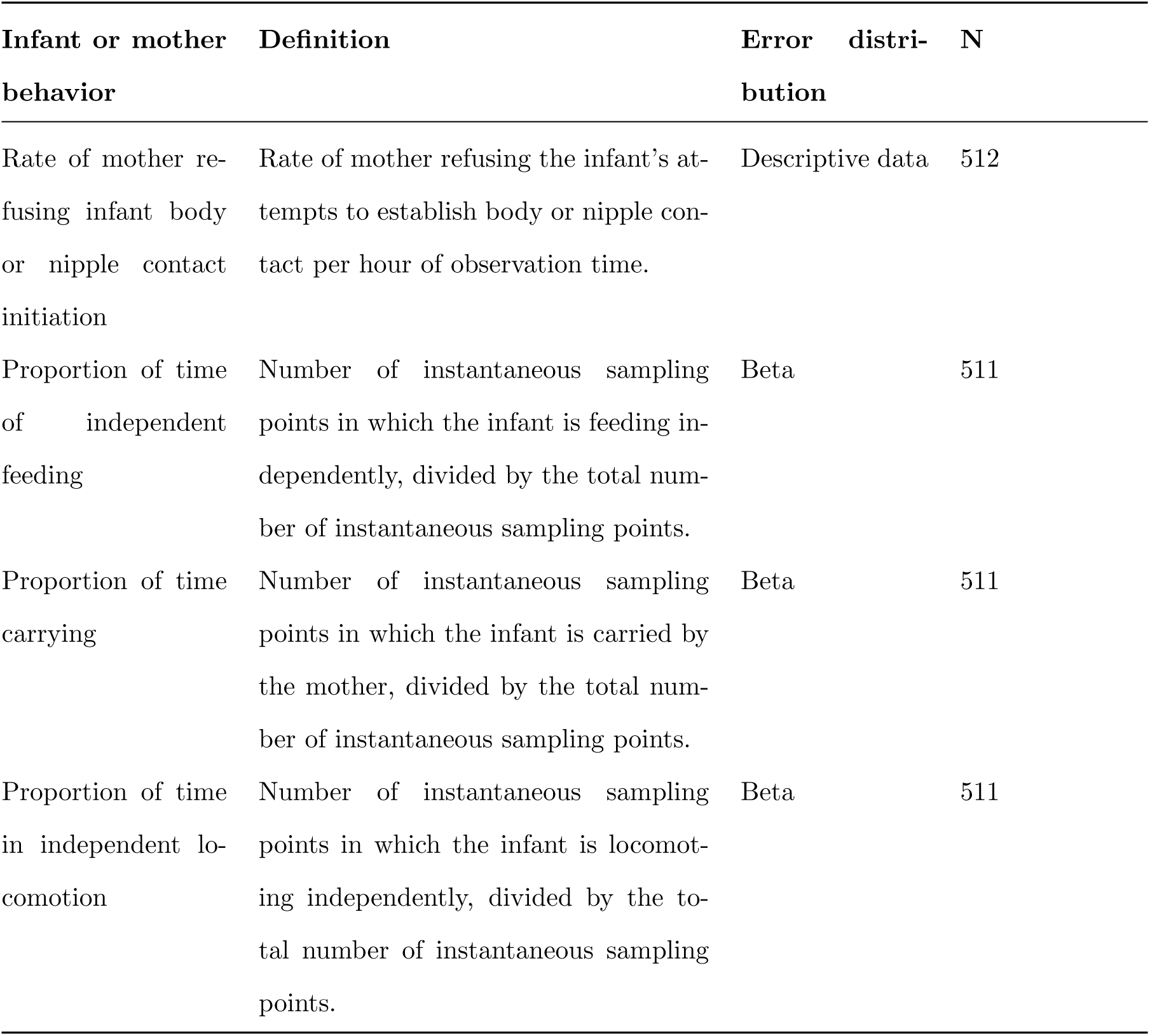
Infant and mother measures analyzed.

To describe the age-trajectories of the mother-infant spatial relationship and the transition from dependent to independent feeding and locomotion, we extracted rates, proportions, or indices based on the infant and mother behaviors (Table S1). These measures have been traditionally used in the literature to study mother-infant relationships (Table 1) and constituted the response variables in the fitted models. Given the scarcity of events observed during the data collection period, we could not fit models for the rate of restraints and for the rate of mother refusing infant body or nipple contact initiation, but we provide some descriptive statistics for these two variables.

We fitted a series of nonlinear generalized mixed models using the Bayesian software package brms (version 2.22.0, Bürkner, 2023) in R version 4.4.0 (R Core Team, 2024). These models incorporated a binomial error distribution for discrete proportions and a beta error structure for continuous proportions, both combined with an identity link function (see Table 1). The models included the Gompertz as described above. To estimate the parameters *c* and *b* in an unconstrained space while constraining their effective parameters to be positive (as they represent scaling factors that cannot be negative), we exponentiated them in the model fitting function. Similarly, the effective parameters *a* and *d* needed to be constrained within the range between 0 and 1, but we wanted to estimate them in an unconstrained space. Hence, we applied an inverse logit transformation in the model fitting function (e.g., 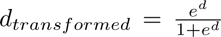). Additionally, to ensured that *d* + *a* does not exceed 1, since each response variable is bound between 0 and 1, we multiplied the effective value of *a* by the effective value of *d*. Thus, the amplitude of change was smaller than the initial asymptote ensuring that the final asymptote was larger than or equal to 0. Two exceptions were the models for the proportion of independent locomotion and feeding time, for which the amplitude of change was larger than the initial asymptote, and hence we multiplied the effective value of *a* by 1 - effective value of *d*. Since the way we constrained the effective parameters *d* + *a* ensured that the fitted model bound remained between 0 and 1, we could, and had to, fit all the models with identity link function.

Additionally, we included fixed effects for fruit availability, maternal parity, and maternal dominance rank, as described above. To simplify the interpretation of the model estimates, fruit availability was standardized using z-scores (with a mean of 0 and a standard deviation of 1) (Schielzeth, 2010). Parity and infant sex were added as binary dummy coded categorical variables. For infant sex, ”female” was designated as the reference level (intercept). For parity, ”multiparous” was defined as the reference level. Infant sex was initially included as an interacting term with infant age, by adding a coefficient for infant sex to each parameter of the Gompertz function described in Equation 1 (*ds*, *as*, *bs* and *cs*). The fixed part of the model formula implemented in brms describing the relationship between the mother and infant behaviors (*y*) and infant age (*x*) is provided in the Supplementary material (Equation 1).

Since no differences across age were found between sexes (Figure S2), we reduced the models by only adding independent effects for infant sex and age. Given that we had repeated measurements for infant, mother, and group ID, both initial and reduced models included them as random intercepts. Since age and fruit availability varied within each mother, infant, and group ID, these variables were also included as random slopes within each grouping factor (Schielzeth & Forstmeier, 2009). We included a random slope for each of the four parameters of infant age (*d*, *a*, *b* and *c*) within all three grouping factors and for each interactive term of infant age and sex (*ds*, *as*, *bs* and *cs*) within mother ID and group ID. Finally, for group ID, we specified infant sex, maternal parity, and maternal rank as additional random slopes.

We chose weakly informative and normally distributed priors for the fixed effects (Table S3). In the case of the parameters of the Gompertz function, we chose their means based on a visual inspection of the data, with a standard deviation of 1 for parameters *a*, *c* and *d*, and a standard deviation of 2 for parameter *b*. For all other fixed effects we chose weak priors, namely a normal distribution with a mean of 0 and a standard deviation of 2. For the prior distributions of standard deviations of the random effects, we used weakly regularizing exponential priors with a rate of 1. We ran 10 000 iterations for each of 8 chains in the MCMC sampling process, with an adapt delta parameter set to 0.85. The Effective Sample Size (ESS) values for Bulk ESS and Tail ESS were greater than 4 000 for most models, indicating adequate sampling. Exceptions were the models for the mother’s responsibility for maintaining body contact, and the proportion of independent locomotion and feeding time, in which Bulk ESS and Tail ESS values were higher than 1 000. Additionally, all models showed R-hat values of 1, indicating no convergence issues, except for the proportion of carrying time model. For this model, R-hat values reached up to 1.06. To address this, we increased the adapt delta parameter to 0.99 and relaxed the prior standard deviations for the *a* and *d* parameters to 2. These adjustments reduced R-hat values to below 1.03 and resulted in Bulk and Tail ESS values exceeding 8000 for fixed effects and 200 for random effects.

For each model, we specified the mean of the priors as the starting values for the parameters to provide stable initialization for the MCMC chains. Posterior predictive checks perform with the Bayesplot R package (Gabry et al., 2019) further suggested that the function was appropriate in capturing the age trajectory for most of the variables analyzed, although predicted values were far from observed values for some age ranges (Figure S1). Also, the model did not adequately fit the response variable for certain variables, such as the mother’s responsibility in maintaining body contact and proximity. To extract the models’ predictions with regard to age and to quantify the uncertainty in them, we determined fitted values with respect to infant age for all posterior samples of model parameters. These posterior distributions of fitted values were then used to determine the posterior median and its 2.5% and 97.5% credible interval, respectively of the fitted model. Additionally, for each model, we report the location of the inflection point <u>^log(^</u>*<u>^b^</u>*<u>^)^</u>, the initial asymptote (*d*), and the final asymptote (*a*), all expressed on the natural scale after back-transforming them to their constrained space.

Additionally, using the Bayesplot R package (Gabry et al., 2019), we created uncertainty plots that display the median and 95% credible intervals of the random effects, extracted from the posterior samples of each model. The models revealed substantial variation (i.e., relatively large estimated standard deviations for the random intercepts and slopes) as compared to the magnitude of the fixed effects estimates, but the estimated standard deviations were highly uncertain, as indicated by the wide credible intervals (Figure S3). To further investigate the consistency of maternal behavior across mothers (Fairbanks, 1996), we visually compared the fitted values across infant age of mothers who had more than one offspring in the sample and the average age trajectory (*N* = 15; see Figure S4). Although some consistency was suggested by the lack of intersection in fitted values between certain mothers, in most cases, the age curves intersected, and confidence intervals overlapped, indicating a high degree of uncertainty in the estimated mother-specific trajectories. This was expected given that infant age was added as a random slope in the models (Mundry et al., 2023).

## Results

### Effect of sex on the mother-infant relationship

The credible intervals of the infant sex coefficients in most of the reduced models did not include 0, but were very wide (Tables S4 and S5). We visually explored potential sex differences for models in which the lower bound of the credible intervals was higher than 1 (proportion of contact time and proportion of carrying time), which also suggested no notable differences in the response variables with regard to infant sex (Figure 1). Consequently, we do not present sex-differented results in the subsequent analyses.

**Fig. 1.**
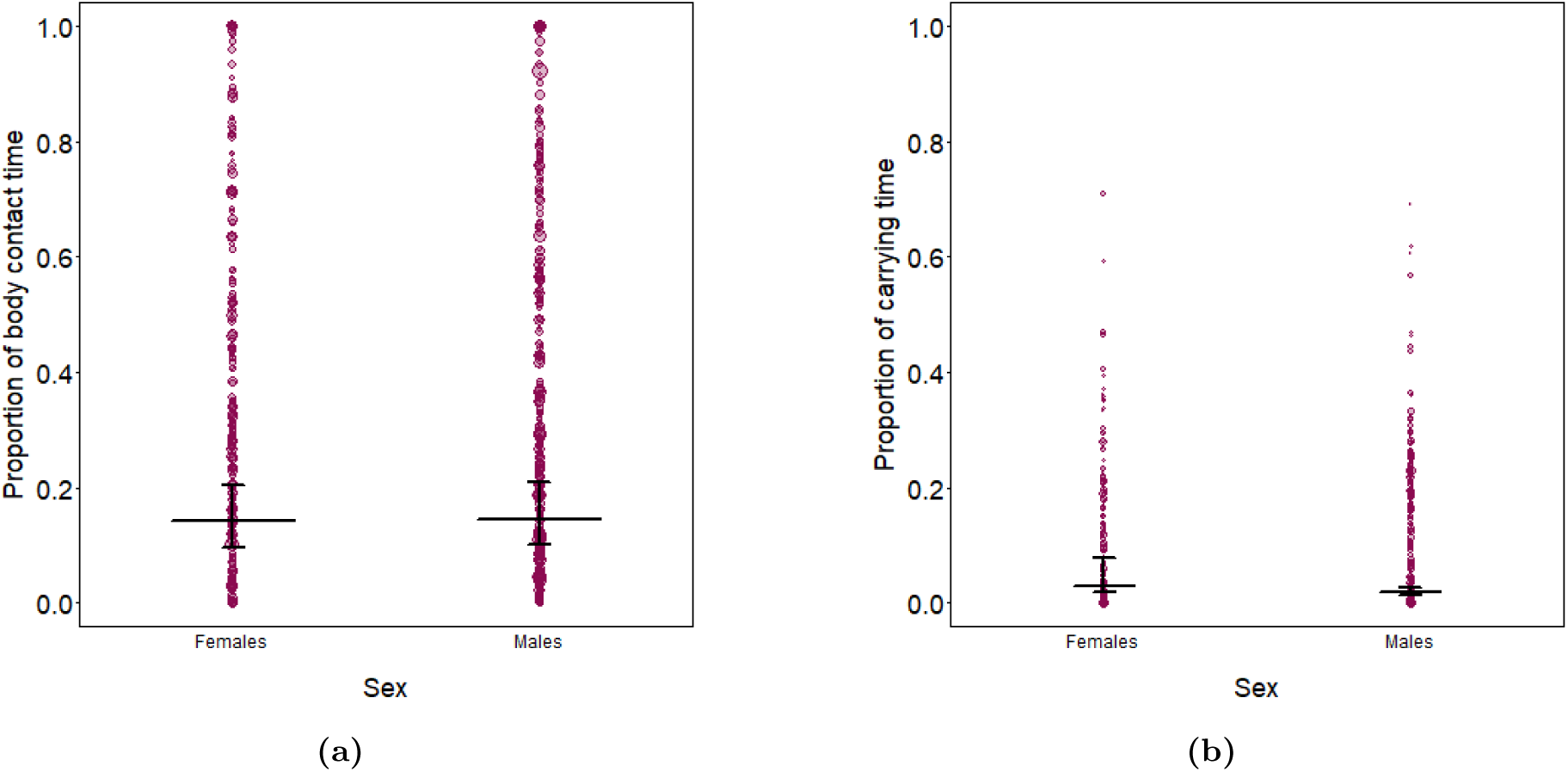
Effect of sex on the (a) proportion of contact time and (b) proportion of carrying time. Dots represent block/infant data points, with the area of each dot proportional to the observation effort for that data point (proportion of contact time: *min*=1.06 h, *max* =16.05 h; proportion of carrying time: *max* =33 sampling points, max=505 sampling points). The horizontal lines indicate the median (50th percentile) of the posterior samples, while the vertical lines represent the 95% credible intervals

### Age-trajectory of the mother-infant spatial relationship

During the first weeks of an infant’s life, mothers spent the majority of the observation time within 1.5m (Figure 2a) and in body contact (Figure 2b) with their infants. Both responses followed a decrease during the subsequent months with the inflection point occurring around 2 months of age. Starting from month 7, the predicted proximity and body contact time fell below 25% (Table 2).

**Fig. 2.**
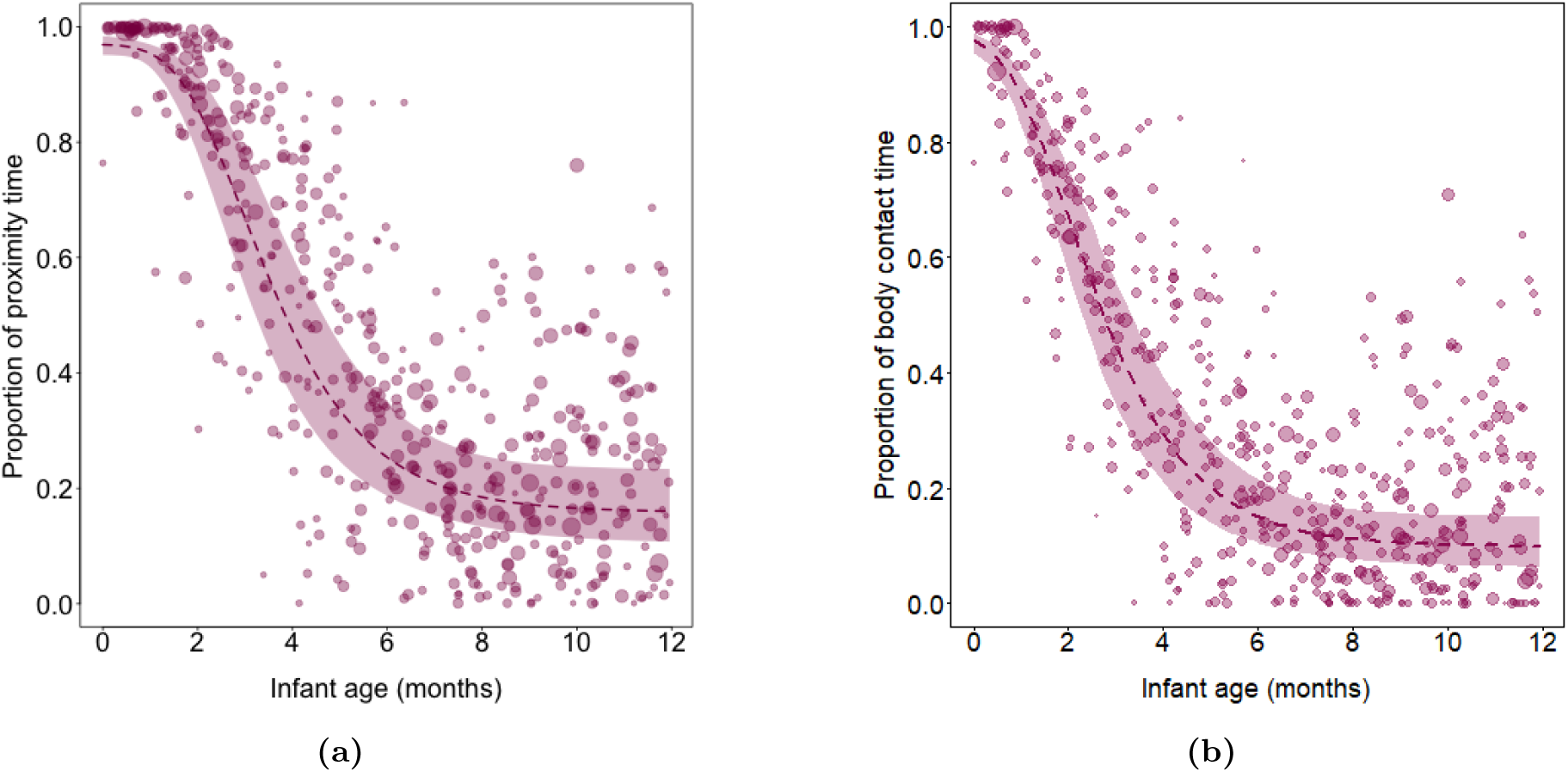
Effect of infant age on the proportion of time in mother-infant (a) proximity and (b) body contact. Dots represent block/infant data points, with the area of each dot proportional to the observation effort for that data point (*min*=1.06 h, *max* =16.05 h). The dashed line represents the median (50th percentile) of the posterior samples at each age value, while the shaded region corresponds to the 95% credible interval of the model predictions, determined as the 2.5th and 97.5th percentiles of the posterior samples Mothers were primarily responsible for maintaining body contact and proximity with their offspring during the first months of infancy (Figures 3a and 3b), with initial asymptotes above 0.5 (see Table 2). This responsibility shifted to the infant when it reached 2 to 4 months of age.

**Table 2.** Inflection point and initial and final asymptote for mother-infant spatial relationship models.

| Model | Inflection point [95% CI] | Initial asymptote [95% CI] | Final asymptote [95% CI] |
| --- | --- | --- | --- |
| Proportion of proximity time | 3.0 age-months [2.5, 3.6] | 0.97 [0.95, 0.98] | 0.15 [0.08, 0.24] |
| Proportion of body contact time | 2.0 age-months [1.7, 2.5] | 0.99 [0.96, 0.99] | 0.10 [0.06, 0.16] |
| Mother responsibility in maintaining proximity | 2.5 age-months [1.6, 3.5] | 0.59 [0.48, 0.76] | 0.30 [0.10, 0.60] |
| Mother responsibility in maintaining body contact | 2.5 age-months [1.5, 4.0] | 0.84 [0.70, 0.95] | 0.30 [0.10, 0.50] |
| Proportion of mother proximity initiation | 2.3 age-months [1.5, 3.4] | 0.55 [0.46, 0.65] | 0.31 [0.21, 0.46] |
| Proportion of infant proximity initiation | 2.4 age-months [1.5, 3.7] | 0.52 [0.45, 0.58] | 0.72 [0.66, 0.79] |
| Proportion of mother body contact initiation | 3.1 age-months [2.6, 3.8] | 0.87 [0.83, 0.92] | 0.32 [0.19, 0.49] |
| Proportion of infant body contact initiation | 3.6 age-months [2.5, 4.8] | 0.38 [0.36, 0.42] | 0.69 [0.61, 0.76] |

**Fig. 3.**
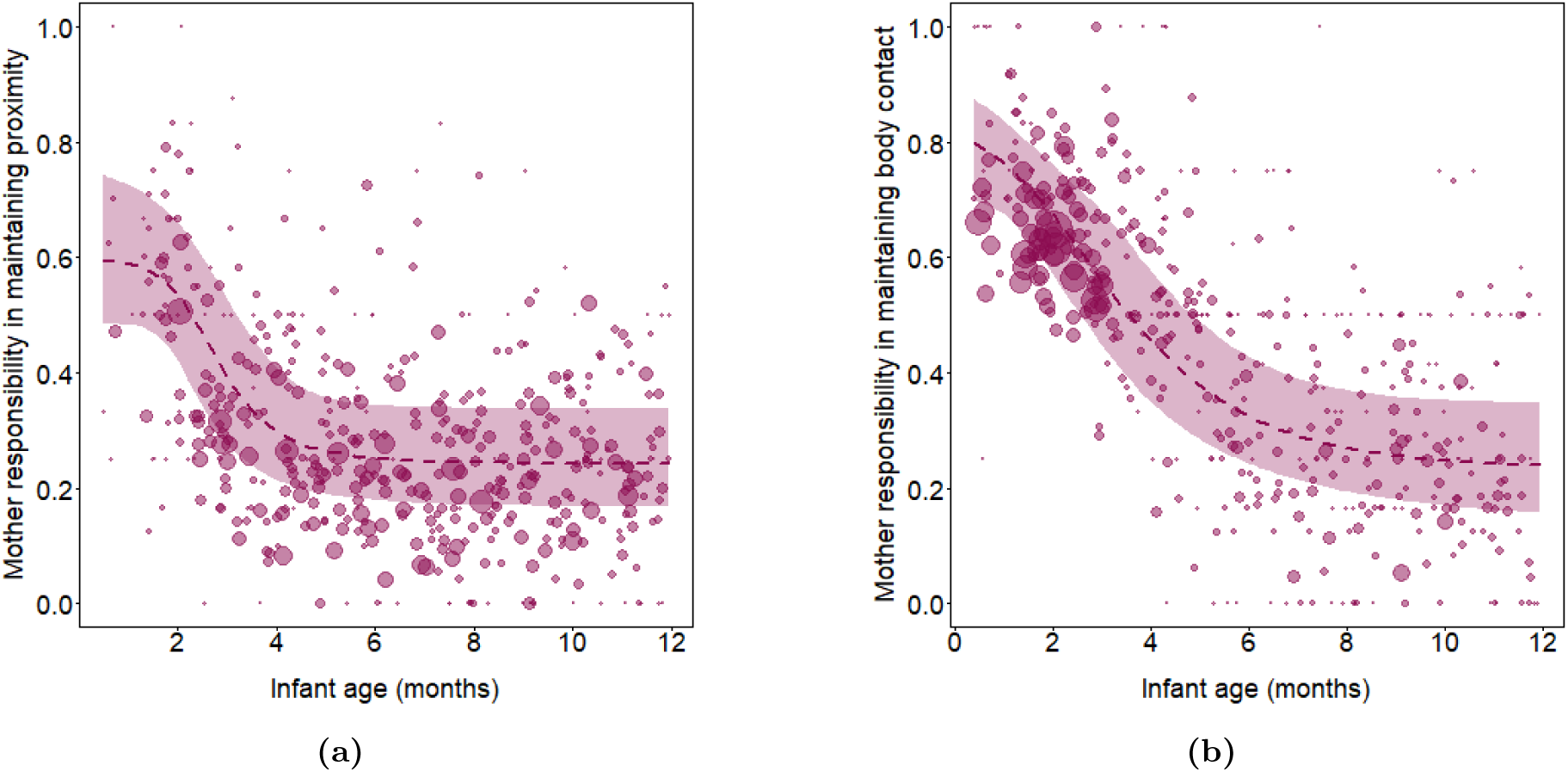
Effects of infant age on the mothers’ responsibility in maintaining (a) proximity and (b) body contact. Dots represent block/infant data points, with the area of each dot proportional to the observation effort for that data point (*min*=2, *max* =366). The dashed lines represent the median (50th percentile) of the posterior samples at each age value, while the shaded region corresponds to the 95% credible interval of the model predictions, determined as the 2.5th and 97.5th percentiles of the posterior samples

The number and area of the dots in Figures 4a and 4b indicate that at the beginning of infancy, there were very few changes in proximity between mothers and infants, whereas the opposite was observed for body contact initiation (Figures 4c and 4d). During the initial months, mothers initiated more interactions than they terminated, both in terms of proximity and body contact (Table 2). However, this pattern gradually shifted, with mothers terminating more interactions than they initiated as the infant grew older. As shown in Figures 4a and 4b, and according to the inflection points, the shift occurred between months 2 to 4.

**Fig. 4.**
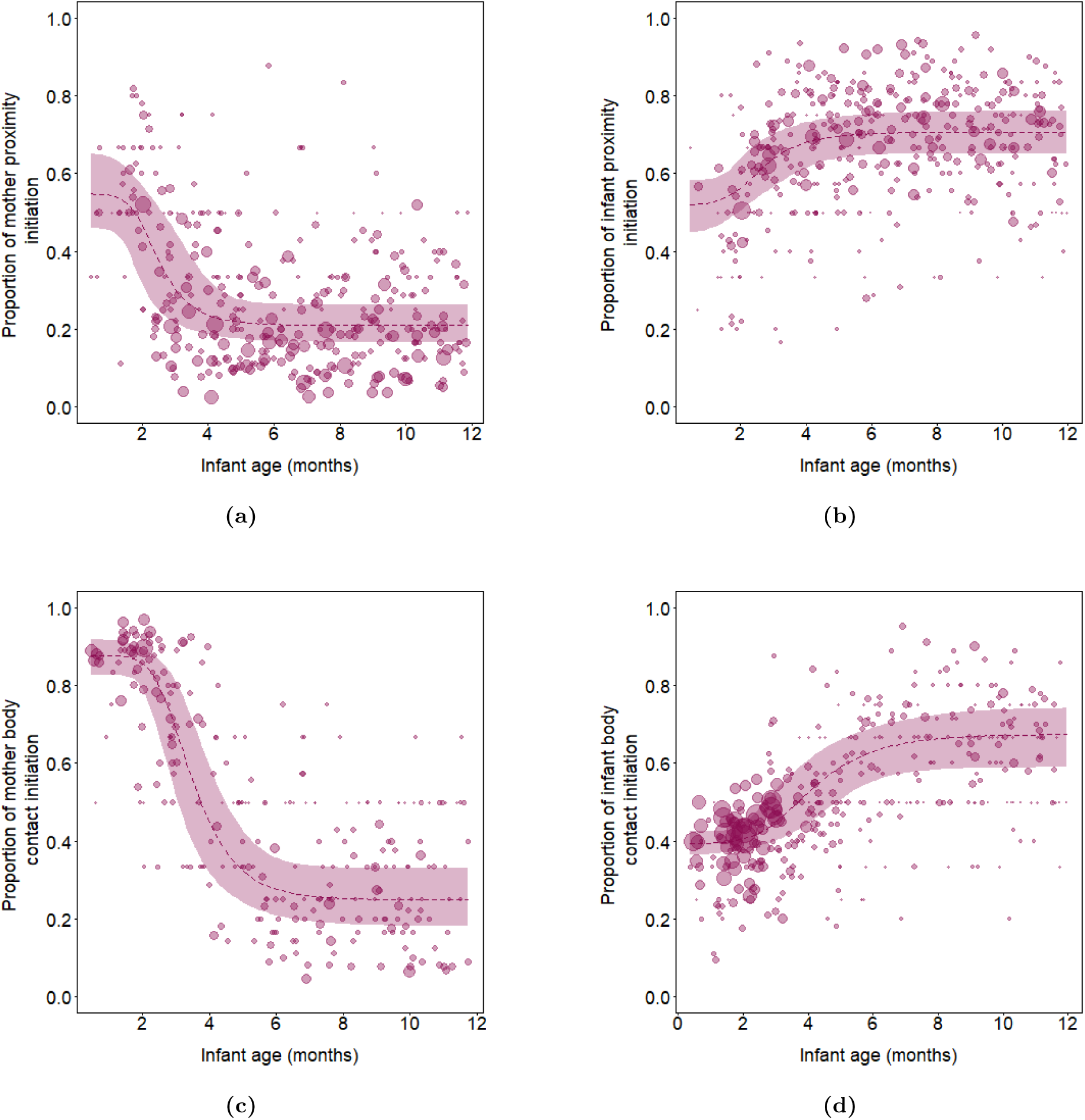
Effect of infant age on the proportion of (a) mother proximity initiation, (b) infant proximity initiation, (c) mother body contact initiation, and (d) infant body contact initiation. Dots represent block/infant data points, with the area of each dot proportional to the observation effort for that data point (*min*=2, *max* =299). The dashed line represents the median (50th percentile) of the posterior samples at each age value, while the shaded region corresponds to the 95% credible interval of the model predictions, determined as the 2.5th and 97.5th percentiles of the posterior samples *max* =8.89).

The pattern described above was reversed for the proportion of infant body contact initiation, which increased by about 20% from month 1 to 12 (Figure 4d and Table 2). In this case, fitted values were above 0.5 from around month 4. Although the proportion of infant proximity initiation also increased from the beginning of infancy to the end (Figure 4d and Table 2), infants consistently approached their mothers more frequently than they left them throughout infancy.

Restraining events were registered only in 24.6% of the data points (*n*=744 events). The time window of restraining started with an infant of 3 days of age and ended at 10.2 months of age. For the subset of data points in which restraint occurred, the mean age was 2.3 months (*sd* =1.5). When considering all data points, the average restraining rate was 0.33 events per hour(*sd* =1.01, min=0,

### Age-trajectory of the transition from dependent to independent feeding and locomotion

Nipple contact time accounted for around 40% of the observation time in newborns, gradually decreasing to less than 10% by month 7, and varied considerably between individual observation blocks (Figure 5a and Table 3). With respect to carrying time, mothers carried their newborns around 20% of the observation time, with fitted values remaining above 20% until month 2 (Figure 5b and Table 3). By around 4 months of age, mothers carried infants for 10% or less of the observed time, and after 7 months of age, the majority of individual observation blocks had no occurrence of maternal carrying, but carrying was still observed twice in an infant of 11.8 months of age.

**Fig. 5.**
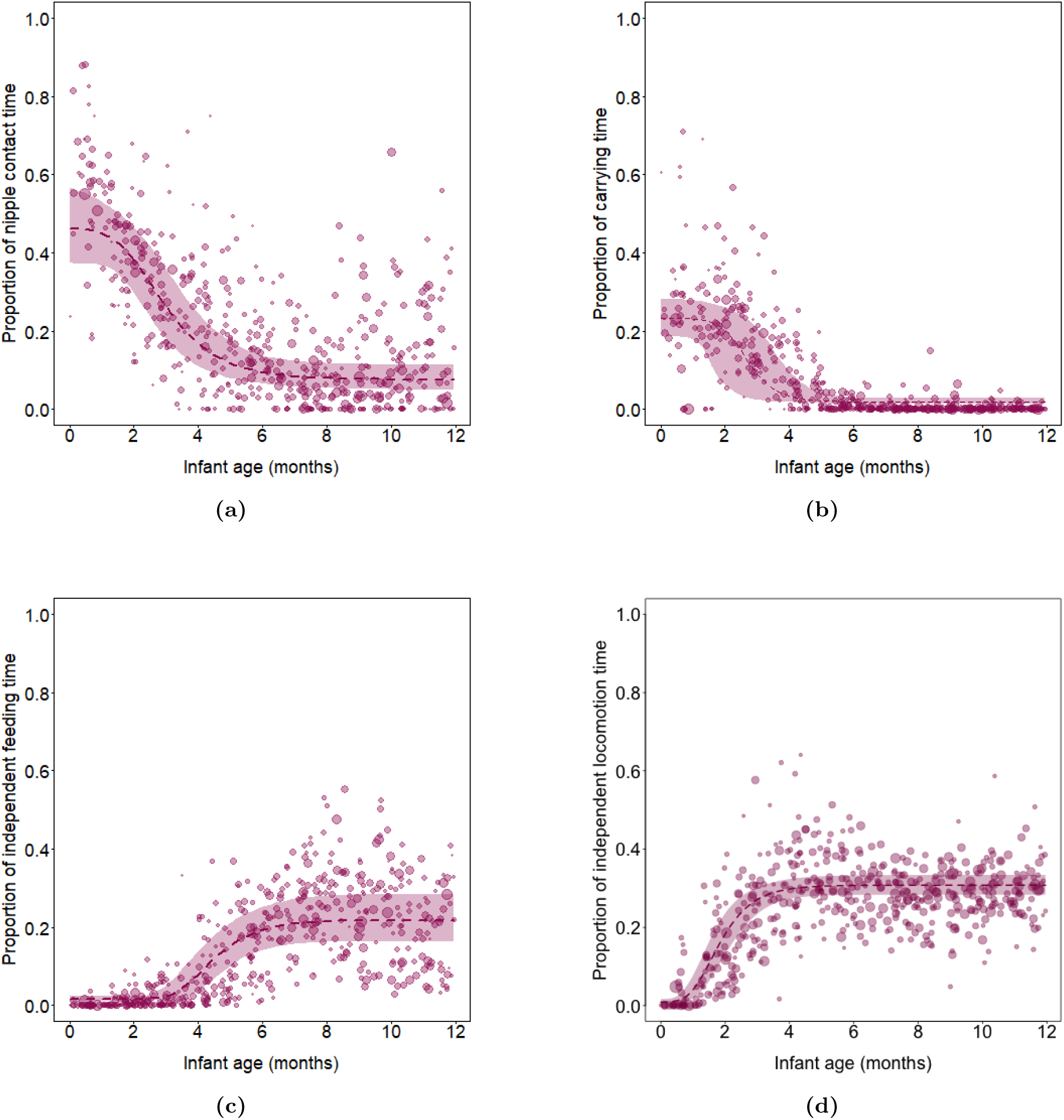
Effect of infant age on the proportion of time in (a) nipple contact, (b) carrying, (c) independent feeding, and (d) independent locomotion. Dots represent block/infant data points, with the area of each dot proportional to the observation effort for that data point (*min*=33 sampling points, *max* =505 sampling points). The dashed lines represent the median (50th percentile) of the posterior samples at each age value, while the shaded regions correspond to the 95% credible interval of the model predictions, determined as the 2.5th and 97.5th percentiles of the posterior samples

**Table 3.** Inflection point as well as initial and final asymptote for the transition from dependent to independent feeding and locomotion models.

| Model | Inflection point [95% CI] | Initial asymptote [95% CI] | Final asymptote [95% CI] |
| --- | --- | --- | --- |
| Proportion of nipple contact time | 2.5 age-months [1.7, 3.4] | 0.46 [0.37, 0.57] | 0.10 [0.10, 0.10] |
| Proportion of independent feeding time | 4.1 age-months [3.5, 4.7] | 0.06 [0.03, 0.11] | 0.43 [0.32, 0.56] |
| Proportion of carrying time | 2.4 age-months [1.3, 3.5] | 0.23 [0.18, 0.28] | 0.01 [0.01, 0.02] |
| Proportion of independent locomotion time | 2.1 age-months [1.7, 2.5] | 0.01 [0, 0.02] | 0.61 [0.55, 0.02] |

Infants rarely fed (Figure 5c and Table 3) or locomoted (Figure 5d and Table 3) independently of their mothers during their first month of life, after which they steadily increased their proportion of independent locomotion time until around 3 months of age. In the case of independent feeding, a steady increase was observed starting around 3 months of age until around 7 months of age.

We observed a total of 356 events of mothers refusing infant body or nipple contact initiation (in 32.0% of the individual observation blocks), distributed across almost all infancy (first case observed at 1.4 months of age and last case at 11.8 months of age). The mean age for the subset of the data which comprised the observed cases of mothers refusal was 7.2 months of age (sd=2.7). In the full data set, mothers refused their infants an average of 0.16 times per hour (sd=0.35, min=0, max= 2.7).

## Discussion

Our findings align with the age-trajectory of mother-infant relationships commonly described in primates and support the general predictions outlined in the Introduction. At birth, the infants of our population of Assamese macaques were almost fully dependent on their mothers, who displayed substantial investment in their offspring in terms of feeding and transportation. A transitional phase emerged early, between 1 and 3 months of age, marked by a noticeable reduction in maternal care. This coincided with a significant increase in independent locomotion and with the start of independent feeding by infants. By 6 to 7 months of infant age, the rate of decrease in maternal care and the rate of increase in infant independence had slowed down. Specifically, body contact and proximity between mothers and infants had decreased substantially (from 100% to less than 50% of the observed time in most data points), and infants had achieved near-complete independence in locomotion. Although nipple contact persisted (as observed in previous studies; Berghäanel et al., 2016), infants spent nearly twice as much time feeding independently. Finally, contrary to our expectations, the age-trajectory and the average levels of maternal care and infant independence across infancy did not change depending on the infant sex, although the high uncertainty of the sex effect in all the models, probably related to the limited sample size of the study, warrants cautious interpretation. The lack of sex differences might suggest, however, that such differences depend on other factors, such as mother age (Soben et al., 2023), or that they emerge later in development, particularly during the juvenile period (Kulik et al., 2016), which corresponds with the increasing sex differences in body size observed with increasing individual age in our population (Anzàa et al., 2022).

Similar to other macaque species (Berman, 1980; Dittus & Baker, 2024; Lindburg, 1971), by the second half of the first year of the infant’s life, maternal care was significantly reduced and correspondingly infants were highly independent. As commonly observed in primates (Fairbanks, 1996; R. A. Hinde & Spencer-Booth, 1967; Lindburg, 1971), the push toward infant independence was largely initiated by mothers, with infants adapting accordingly. During the initial months, infants were highly active, frequently breaking physical contact with their mothers to explore their surroundings. Mothers, in turn, actively sought to maintain contact, often restraining their infants’ attempts to move away. However, in the transitional phase, mothers began to refuse contact attempts and increased spatial distance from their offspring, while infants responded by intensifying efforts to maintain proximity and re-establish contact. This pattern lead to older infants being mainly responsible for the remaining spatial proximity with their mothers.

As in other primates, locomotor independence preceded feeding independence: a gap of around 2 months with respect to the first signs of independence, and a larger gap in the full achievement of independence (carrying ended around 6 months of age, while nursing persisted until the end of infancy). Earlier cessation of carrying compared to nursing has been explained by the higher energetic costs of the former (Altmann & Samuels, 1992; Young & Shapiro, 2018). The timing of the cessation of carrying in our population was similar to wild toque macaques (*Macaca sinica*) (Dittus & Baker, 2024) and free-ranging rhesus macaques (Lindburg, 1971). From the infants’ perspective, the early initiation of independent locomotion described in our study, also corresponds with free-ranging rhesus macaque infants starting to locomote during the second month of age (Dittus & Baker, 2024; Lindburg, 1971). This might be related to all major locomotory skills being already acquired by that age (Berghäanel et al., 2015).

Regarding the other milestone in mammalian infant independence, age of weaning, two events must be considered: the end of the mother’s milk provision (Eisenberg, 1993), and the first day in which infants eat solid food (Akre, 1990). Our infants began to consume solid food as early as 2 months, which is consistent with the mean age reported across 16 cercopithecid species (Langer, 2003), and with observations from other macaque populations (Dittus & Baker, 2024). The start of solid food consumption was accompanied by a significant decrease in nipple contact time during the first half of infancy, coinciding with a peak in maternal rejection at 7 months of age that resembled that of other wild macaques (*Macaca sinica*, Dittus and Baker, 2024; *Macaca silenus*, Krishna et al., 2008; *Macaca thibetana* Deng and Zhao, 1991). It also aligned with a reported average end of lactation in cercopithecines at around 7 months of age (Langer, 2003). However, the infants of our population stayed in nipple contact with their mothers approximately 10% of the observed time, and in some cases above 60%, well into the second half of their first year of life. Milk consumption is difficult to estimate from observation only (K. Hinde, 2009), given that it is often methodologically unfeasible to distinguish between suckling and nipple contact in wild, arboreal populations. Thus, we would need to directly measure the presence or absence of milk consumption to be able to conclude the exact age in which immature Assamese macaques become nutritionally independent of their mothers (for instance, with stable isotope analysis: Crowley and Hinde, 2012). Notably, by directly assessing the mammary tissue of wild toque macaque mothers, Dittus and Baker 2024 reported that while some infants stopped accessing maternal milk by 7.2 months, most have available milk until 18 months or later. This suggests that female macaques may employ a mixed feeding strategy to optimize both infant survival and maternal reproductive success. Accordingly, the persistence of nipple contact likely reflects extended energetic maternal support rather than merely comfort, as some studies have proposed (e.g., Băadescu et al., 2016). Nonetheless, the high interindividual variation, and the lacking information about nocturnal suckling events, demand further investigation of the weaning transition in our population of Assamese macaques.

Through a prolonged mixed-feeding strategy, Assamese macaque mothers reduce their energetic investment by dividing infants’ intake between maternal milk and foraged solid food (Langer, 2003). As seasonal breeders, Assamese macaques have evolved a relaxed income breeding strategy in response to fluctuating energy availability in their environment (Brockman & van Schaik, 2005; Touitou et al., 2021). Specifically, lactation peaks are synchronized with periods of high food availability, supporting early infant survival. Females also build fat stores during late lactation to facilitate future conception (Heesen et al., 2013; Touitou et al., 2021), and their probability of conceiving increases with better physical condition (Heesen et al., 2013). Given their seasonal reproductive pattern (Fürtbauer et al., 2010), mothers following a 1-year interbirth interval must shift energy allocation to future offspring during the second half of the current offspring’s infancy. Although this shift significantly reduces infant survival (Shivani et al., 2025), particularly during late infancy, it may be partially compensated by the prolonged mixed-feeding period. Further research with larger sample sizes should explore the impact of interbirth intervals on the mother-infant relationship, as seen in chacma baboons (*Papio ursinus*), in which infant tantrums were linked to maternal reproductive strategies (Dezeure et al., 2021). This question can only be answered in wild populations, in which limited food availability forces females to allocate energy to one reproductive process at the expense of another.

The Gompertz function has proven a valuable tool to describe how the mother-infant relationship in a wild primate population changes as a function of infant age. While it provided a robust fit for many response variables, it did not perform well for others. Further exploration of alternative nonlinear functions, such as other sigmoidal functions (Zullinger et al., 1984), may improve model fit and parameter interpretation. Although generalized additive mixed models (GAMMs) are a potential alternative, we opted for the Gompertz function to avoid the overfitting and biologically unrealistic values that GAMMs can produce. Furthermore, the Gompertz function offers biologically meaningful parameters, such as initial and final asymptotes and inflection points, that can be estimated and for which data can be simulated.

This study provides a precise quantitative description of the age trajectory of the mother-infant relationship in a wild population of Assamese macaques. The analysis of several maternal care and infant independence behaviors allowed us to describe how the mother-infant dyad develops as a unit, with the behavior of both members changing synchronically (Moore, 2007; Provenzi et al., 2018; Rosenblatt, 1965). To further disentangle the development of the behavioral feedback between dyad partners, analysis with sufficient samples sizes should directly asses how the mother-infant relationship emerges moment-to-moment (e.g., time series analysis, Berkhout et al., 2023). The observed transition toward independence aligns with a relaxed income breeding strategy, in which mothers in sufficient physical condition can optimize reproductive output by decreasing maternal care during late infancy and reallocating energy to future offspring. Consequently, infants become primarily responsible for maintaining interactions with their mothers, who continue to provide some maternal support. Future studies with larger sample sizes are needed to assess how individual, group, and population-level variables may impact the mother-infant relationship of our study species. Given the complex relationship between life-history traits and both intrinsic and extrinsic species characteristics, direct comparative studies are necessary to better situate and understand the position of Assamese macaques within the slow-fast life history continuum (e.g., Castellano-Navarro et al., 2023). Finally, our findings underscore the importance of employing non-linear, rather than linear, models to fully capture the continuities and discontinuities in the development of the mother-infant relationship.

## Statements and Declarations

### Ethics approval

Data collection was authorized by the Department of National Parks, Wildlife and Plant Conservation (DNP), and the National Research Council of Thailand (NRCT) under a benefit-sharing agreement (permit numbers: 0002.3/2647, 0002/17, 0002/4137, 0402/2798, 0401/11121). The study was purely observational and followed the ASAB/ABS guidelines for the use of animals in research (https://www. asab.org/ethics).

## Data availability

The data supporting the findings of this study are openly available in GRO.data, under the title Replication data for: ”Age-trajectory of mother-infant relationships in wild Assamese macaques”. The dataset includes raw data and an example R script.

## Acknowledgments

We are grateful to the National Research Council of Thailand (NRCT) and the Department of National Parks, Wildlife and Plant Conservation (DNP) for permission to carry out this research at the Phu Khieo Wildlife Sanctuary. We thank the superintendents of Phu Khieo Wildlife Sanctuary, K. Nitaya, T. Wongsnak, M. Pongjantarasatien, and C. Intumarn, for the support granted and N. Bhumpakphan, Department of Forest Biology, Kasetsart University, for cooperation. We are grateful to Andreas Koenig and Carola Borries at Stony Brook University for developing the field site at Huay Mai Sot Yai. We thank Andreas Berghäanel, Christin Minge, Simone Anźa, and the long-term field team members who contributed to data collection and Andreas Berghäanel and Simone Anźa for fruitful discussions.

