## Supplementary material for "Age-trajectory of mother-infant relationships in wild Assamese macaques"

### Ethogram of mother and infant behaviors

**Table 1** Ethogram of mother and infant behaviors

| Behavior | Definition | Type of sampling |
|---|---|---|
| Proximity initiation | The mother or infant approaches the other member of the dyad within 1.5m (this behavior was always recorded even when the individual only passed by and moved swiftly in and out of the 1.5m sphere) | Continuous |
| Proximity termination | The mother or infant leaves a 1.5m radius around the other member of the dyad | Continuous |
| Body contact initiation | The mother or infant comes so close to the other member of the dyad that parts of their bodies touch (physical contact during fighting was not recorded) | Continuous |
| Body contact termination | The mother or infant increases distance to the other member of the dyad, until no parts of their bodies touch anymore | Continuous |
| Mother restraints | The mother restrains the movement of the infant by holding its arm, leg, foot or tail. This was not coded as a state but as an event (i.e., recorded only once even if the behavior lasted a long period of time). It was recorded again every time the mother stopped holding the infant and started holding it again. | Continuous |
| Mother refuses infant body or nipple contact initiation | The infant tries to make body or nipple contact with the mother and the mother refuses to allow it by pushing the infant away, moving away her nipple with her hand or turning away. It was recorded again every time the mother refused contact, even if it was part of the same behavioural sequence. | Continuous |
| Nipple contact | The infant is in oral contact with the mother's nipple. | Instantaneous |
| Independent feeding | Infant manipulates or ingests food that is not the mother's milk. | Instantaneous |
| Carrying | The infant is on the mother while the mother moves, hangs or stands. | Instantaneous |
| Independent locomotion | The infant walks, runs, trots, jumps or climbs. | Instantaneous |

### Minimum and maximum Elo-ratings within each group-year combination

**Table 2** Minimum and maximum Elo-ratings within each group-year combination

| Group | Year | min Elo-rating | max Elo-rating |
|---|---|---|---|
| stu | 2011 | 542 | 1842 |
| mot | 2018 | 338 | 2201 |
| mst | 2018 | 374 | 2028 |
| sst | 2018 | 440 | 1883 |
| mot | 2022 | 393 | 2281 |
| ms1 | 2022 | 594 | 1487 |
| ms2 | 2022 | 875 | 2323 |
| ms3 | 2022 | 196 | 1050 |
| sst | 2022 | 304 | 1924 |

### Model formula

$$
\begin{aligned}
\texttt{behavior} \sim\ & \frac{\exp(c_D + c_{DS} \cdot \texttt{infant sex})}{1 + \exp(c_D + c_{DS} \cdot \texttt{infant sex})} \\
& - \Bigg( \frac{\exp(c_D + c_{DS} \cdot \texttt{infant sex})}{1 + \exp(c_D + c_{DS} \cdot \texttt{infant sex})} \cdot \frac{\exp(c_A + c_{AS} \cdot \texttt{infant sex})}{1 + \exp(c_A + c_{AS} \cdot \texttt{infant sex})} \\
& \quad \cdot \exp\left(-\exp(c_B + c_{BS} \cdot \texttt{infant sex}) \cdot \exp\left(-\exp(c_C + c_{CS} \cdot \texttt{infant sex}) \cdot \texttt{infant age}\right)\right) \Bigg) \\
& \cdot \frac{\exp(c_{ME} \cdot \texttt{maternal experience} + c_{FA} \cdot \texttt{fruit availability} + c_{MR} \cdot \texttt{maternal rank})}{1 + \exp(c_{ME} \cdot \texttt{maternal experience} + c_{FA} \cdot \texttt{fruit availability} + c_{MR} \cdot \texttt{maternal rank})}
\end{aligned}
\tag{1}
$$

Coefficients in the model formula are represented using the notation $c_X$, where the subscript $X$ corresponds to the associated predictor variable. $a$, $b$, $c$ and $d$ correspond to the parameters of the Gompertz function, $c_{ME}$ denotes the coefficient for maternal experience, $c_{FA}$ the coefficient for fruit availability, and $c_{MR}$ the coefficient for maternal rank.

### Starting values and priors

For the parameters of the Gompertz function, we selected the priors based on visual inspection of the data and used their means as the starting values to provide stable initialization for the MCMC chains. In all models, parameter $b$ had a mean of 2.3 and a standard deviation of 2, while parameter $c$ had a mean of 0 and a standard deviation of 1. The means of parameters $a$ and $d$ varied between models, as shown in Table 3, but each had an standard deviation of 1 across all models.

**Table 3** Means of priors of parameters $a$ and $d$ in all models

| Model | mean prior $d$ | mean prior $a$ |
|---|---|---|
| Proportion of proximity time | 4.6 | 0.85 |
| Proportion of contact time | 4.6 | 1.4 |
| Mother responsibility of maintaining proximity | 0.98 | 0.08 |
| Mother responsibility of maintaining body contact | 2.2 | 0.7 |
| Proportion of mother proximity initiation | 0.6 | 0.85 |
| Proportion of infant proximity initiation | -0.40 | -0.75 |
| Proportion of mother body contact initiation | 2.1 | 0.8 |
| Proportion of infant body contact initiation | -0.84 | 1.3 |
| Proportion of nursing time | 0.98 | 0.08 |
| Proportion of independent feeding time | -5 | -0.8 |
| Proportion of carrying time | -1.38 | -4.5 |
| Proportion of independent locomotion time | -5 | -0.85 |

### Posterior predictive checks

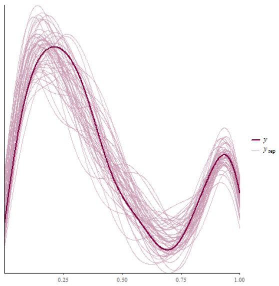

**(a)** Proportion of time in proximity

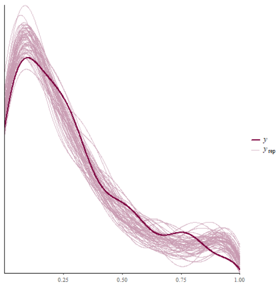

**(b)** Proportion of body contact time

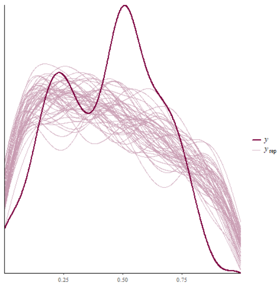

**(c)** Mother responsibility in maintaining proximity

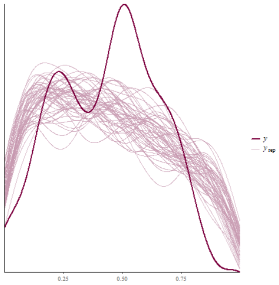

**(d)** Mother responsibility in maintaining body contact

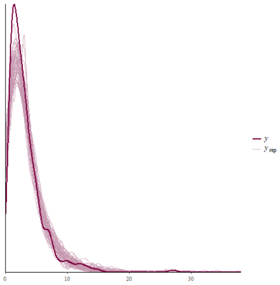

**(e)** Proportion of mother proximity initiation

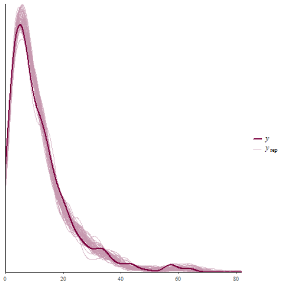

**(f)** Proportion of infant proximity initiation

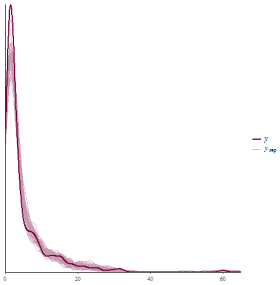

**(g)** Proportion of mother body contact initiation

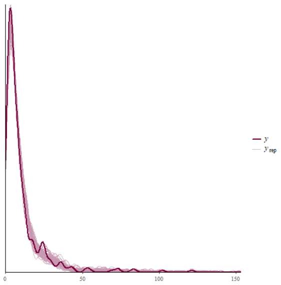

**(h)** Proportion of infant body contact initiation

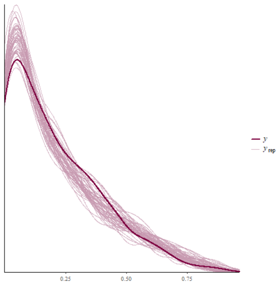

**(i)** Proportion of nipple contact time

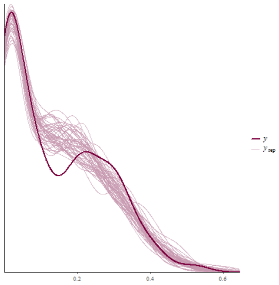

**(j)** Proportion of independent feeding time

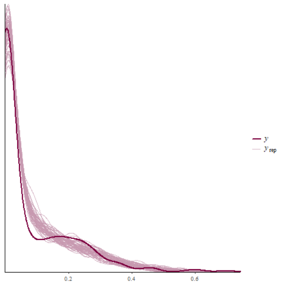

**(k)** Proportion of carrying time

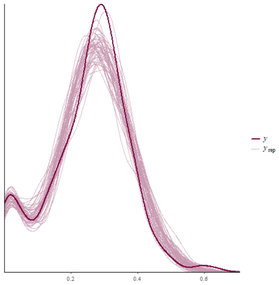

**(l)** Proportion of independent locomotion time

**Fig. 1** Posterior predictive checks performed using the ppc_dens_overlay function from the Bayesplot R package, comparing the observed data with 50 posterior samples.

### Interactive effect of age and sex

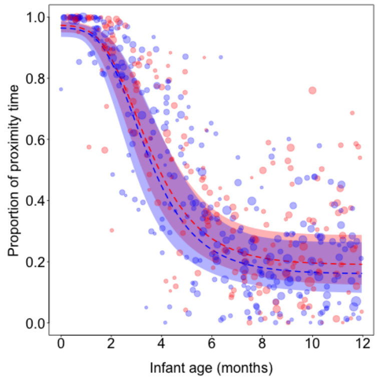


**(a)** Proportion of time in proximity

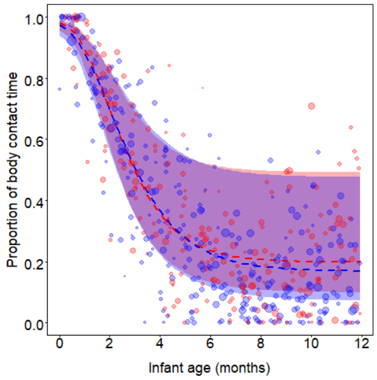


**(b)** Proportion of body contact time

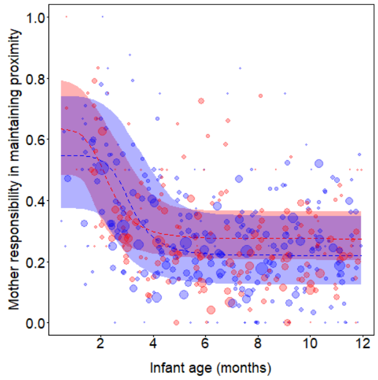


**(c)** Mother responsibility in maintaining proximity

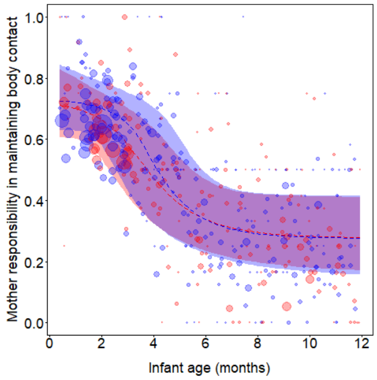


**(d)** Mother responsibility in maintaining body contact

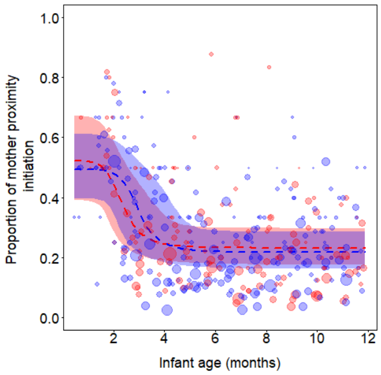


**(e)** Proportion of mother proximity initiation

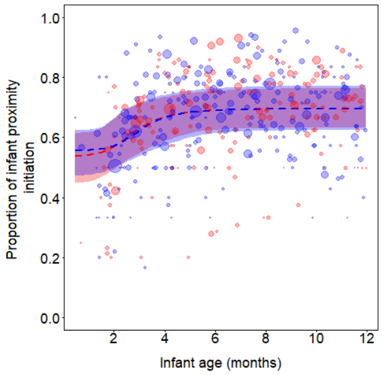


**(f)** Proportion of infant proximity initiation

**Fig. 2** Interactive effects of infant age and sex. Dots represent block/infant data points, with the area of each dot being proportional to the observation effort for that data point (proportion of proximity/contact time: *min*=1.06 h, max=16.05 h; proportion of nipple contact, carrying, independent feeding and independent travel time: *min*=33 sampling points, *max*=505 sampling points; others: *min*=2, max=366). The dashed lines represent the median (50th percentile) of the posterior samples at each age value, while the shaded regions corresponds to the 95% credible interval of the model predictions, determined as the 2.5th and 97.5th percentiles of the posterior samples. Blue dots, lines, and shaded regions are depicting males, and females are depicted in red.

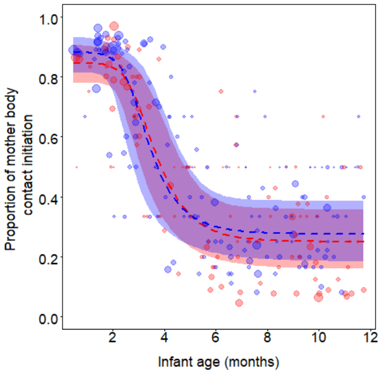


**(g)** Proportion of mother body contact initiation

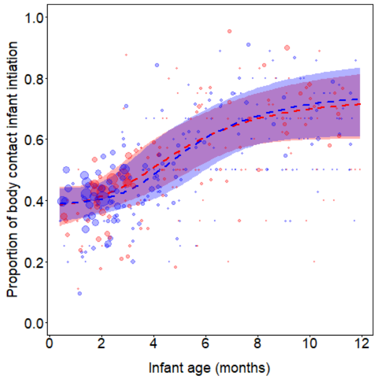


**(h)** Proportion of infant body contact initiation

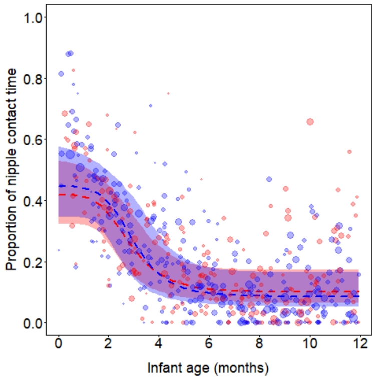


**(i)** Proportion of nipple contact time

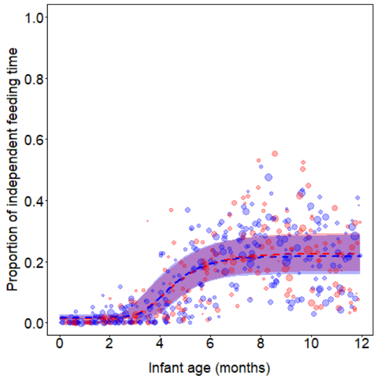


**(j)** Proportion of independent feeding time

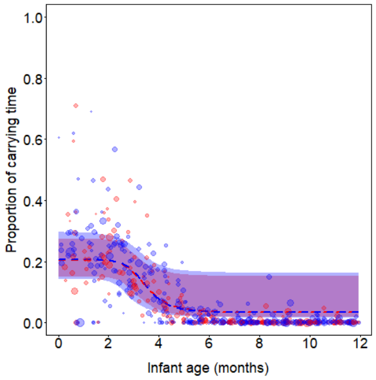


**(k)** Proportion of carrying time

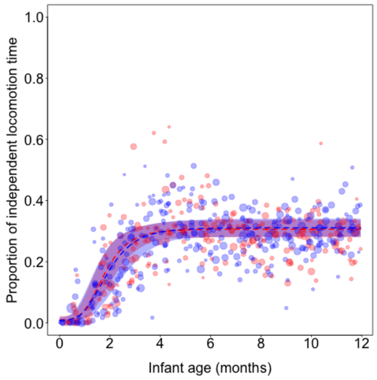


**(l)** Proportion of independent locomotion time

**Fig. 2** Interactive effects of infant age and sex. Dots represent block/infant data points, with the area of each dot being proportional to the observation effort for that data point (proportion of proximity/contact time: $min$=1.06 h, max=16.05 h; proportion of nipple contact, carrying, independent feeding and independent travel time: $min$=33 sampling points, $max$=505 sampling points; others: $min$=2, max=366). The dashed lines represent the median (50th percentile) of the posterior samples at each age value, while the shaded regions corresponds to the 95% credible interval of the model predictions, determined as the 2.5th and 97.5th percentiles of the posterior samples. Blue dots, lines, and shaded regions are depicting males, and females are depicted in red.

### Random variation

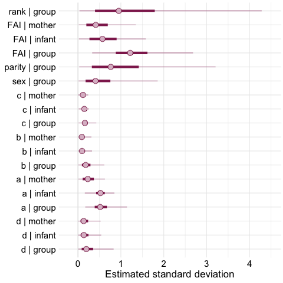


**(a)** Proportion of time in proximity

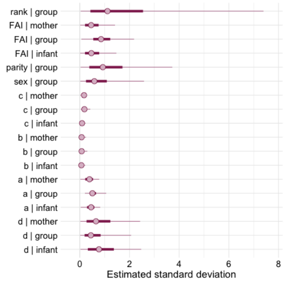


**(b)** Proportion of body contact time

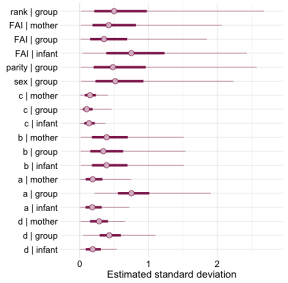


**(c)** Mother responsibility in maintaining proximity

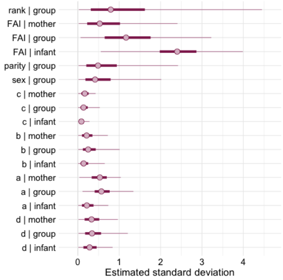


**(d)** Mother responsibility in maintaining body contact

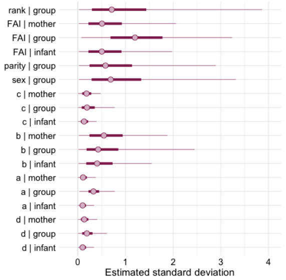


**(e)** Proportion of mother proximity initiation

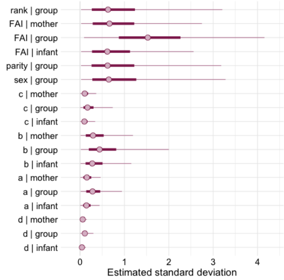


**(f)** Proportion of infant proximity initiation

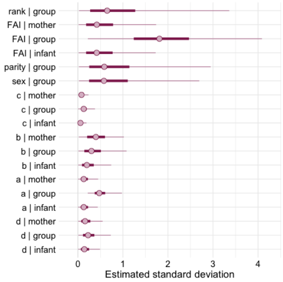


**(g)** Proportion of mother body contact initiation

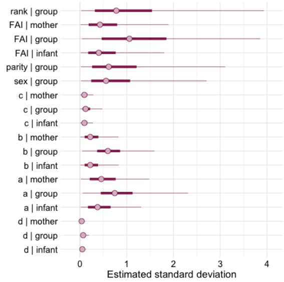


**(h)** Proportion of infant body contact initiation

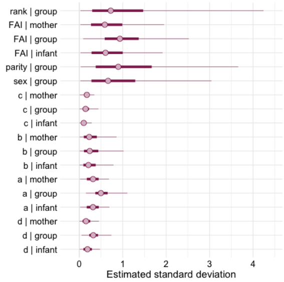


**(i)** Proportion of nipple contact time

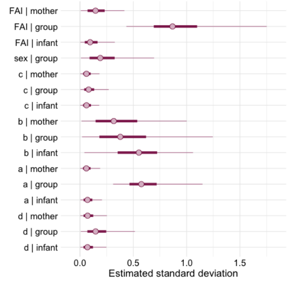


**(j)** Proportion of independent feeding time

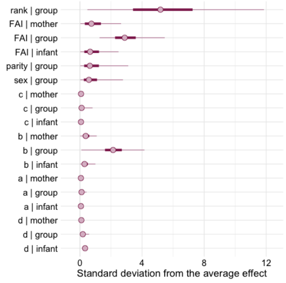


**(k)** Proportion of carrying time

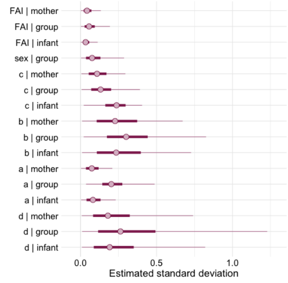


**(l)** Proportion of independent locomotion time

**Fig. 3** Point estimates (50th percentile of the posterior probability distribution) for each random effect, with thin lines capturing 50% of the posterior probability and thick lines representing 95% posterior probability.

### Consistency in mothers with repeated infants

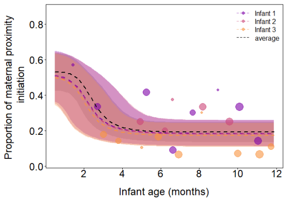


**(a)** Mother 1

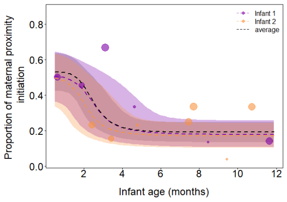


**(b)** Mother 2

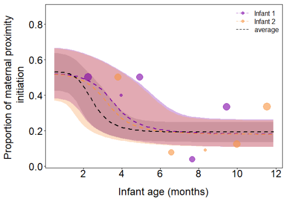


**(c)** Mother 3

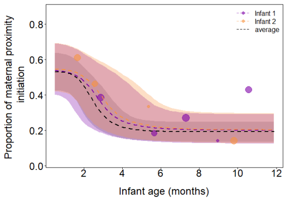


**(d)** Mother 4

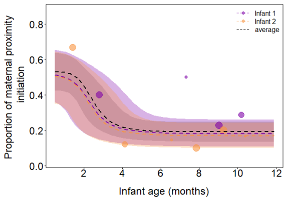


**(e)** Mother 5

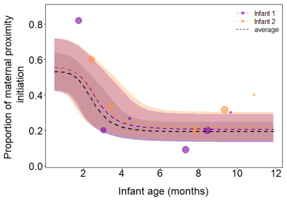


**(f)** Mother 6

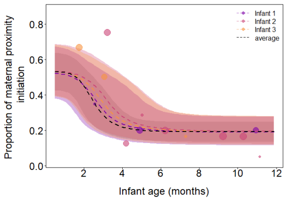


**(g)** Mother 7

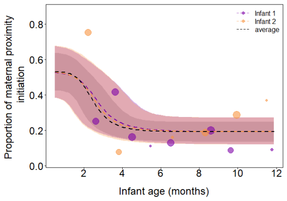


**(h)** Mother 8

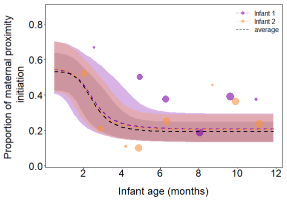


**(i)** Mother 9

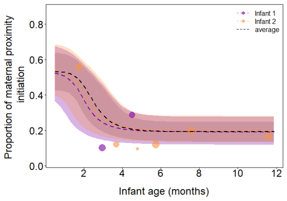


**(j)** Mother 10

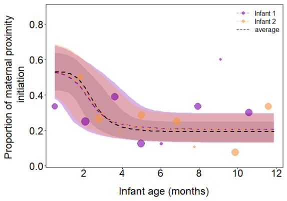


**(k)** Mother 11

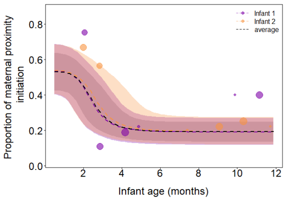


**(l)** Mother 12

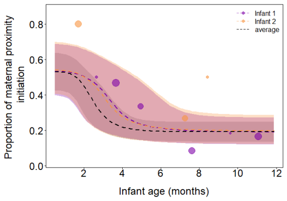


**(m)** Mother 13

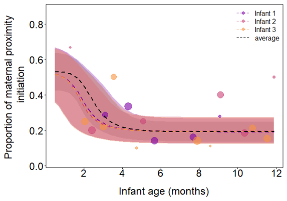


**(n)** Mother 14

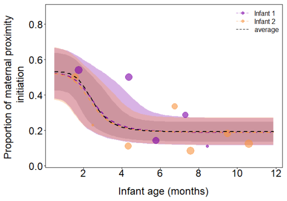


**(o)** Mother 15

**Fig. 4** Age-trajectory of the proportion of proximity initiation for every mother, depicting its average fitted values and the trajectory for each of its infants. Dots represent block/infant data points, with the area of each dot being proportional to the observation effort for that data point *min*=1.06 h, *max*=16.05 h). The dashed line represents the median (50th percentile) of the posterior samples at each age value, while the shaded region corresponds to the 95% credible interval of the model predictions, determined as the 2.5th and 97.5th percentiles of the posterior samples. Black dashed lines represent average fitted values.

#### Regression coefficients for the age-trajectory of the mother-infant spatial relationship

**Table 4** Regression coefficients for the age-trajectory of the mother-infant spatial relationship (mean, standard deviations, and credible intervals of posterior distribution)

| **Coefficient** | **Estimate** | **Standard Error** | **95% CI Lower** | **95% CI Upper** |
|---|---|---|---|---|
| **Proportion of proximity time** | | | | |
| d | 3.49 | 0.31 | 2.97 | 4.18 |
| a | 1.87 | 0.26 | 1.38 | 2.42 |
| b | 2.11 | 0.20 | 1.73 | 2.53 |
| c | -0.35 | 0.12 | -0.58 | -0.12 |
| Sex (male) | 1.65 | 0.74 | 0.50 | 3.37 |
| Parity (nulliparous) | 2.27 | 1.19 | 0.01 | 4.82 |
| Fruit availability [a] | -1.04 | 0.62 | -2.35 | 0.13 |
| Maternal rank | 5.75 | 1.17 | 3.48 | 8.09 |
| **Proportion of contact time** | | | | |
| d | 4.72 | 0.83 | 3.26 | 6.46 |
| a | 2.28 | 0.25 | 1.80 | 2.81 |
| b | 1.47 | 0.11 | 1.26 | 1.71 |
| c | -0.33 | 0.10 | -0.53 | -0.13 |
| Sex (male) | 3.17 | 1.06 | 1.43 | 5.58 |
| Parity (nulliparous) | 2.09 | 1.40 | -0.68 | 4.95 |
| Fruit availability [a] | -0.59 | 0.58 | -1.81 | 0.51 |
| Maternal rank | 6.86 | 1.42 | 3.73 | 9.50 |
| **Mother responsibility of maintaining proximity** | | | | |
| d | 0.43 | 0.31 | -0.06 | 1.17 |
| a | 0.88 | 0.57 | 0.04 | 2.38 |
| b | 3.52 | 1.33 | 1.31 | 6.44 |
| c | 0.28 | 0.34 | -0.39 | 0.91 |

**Table 4 (continued): Response variables**

| Coefficient | Estimate | Standard Error | 95% CI Lower | 95% CI Upper |
|---|---|---|---|---|
| Sex (male) | 1.52 | 0.99 | 0.06 | 3.95 |
| Parity (nulliparous) | 1.59 | 1.29 | -0.37 | 4.59 |
| Fruit availability [a] | -0.04 | 0.53 | -1.05 | 1.14 |
| Maternal rank | 2.20 | 1.29 | 0.14 | 5.04 |
| **Mother responsibility in maintaining body contact** | | | | |
| d | 1.70 | 0.53 | 0.86 | 2.93 |
| a | 1.07 | 0.36 | 0.39 | 1.84 |
| b | 1.49 | 0.68 | 0.65 | 3.40 |
| c | -0.60 | 0.26 | -1.07 | -0.01 |
| Sex (male) | 2.16 | 1.00 | 0.49 | 4.37 |
| Parity (nulliparous) | 0.57 | 1.13 | -1.51 | 2.98 |
| Fruit availability [a] | -0.82 | 0.81 | -2.52 | 0.72 |
| Maternal rank | 5.14 | 1.42 | 2.28 | 7.97 |
| **Proportion of mother proximity initiation** | | | | |
| d | 0.20 | 0.20 | -0.16 | 0.63 |
| a | 0.57 | 0.22 | 0.13 | 0.98 |
| b | 3.66 | 0.76 | 2.23 | 5.22 |
| c | 0.46 | 0.23 | -0.02 | 0.90 |
| Sex (male) | 2.44 | 1.15 | 0.47 | 4.98 |
| Parity (nulliparous) | 1.25 | 1.48 | -1.41 | 4.44 |
| Fruit availability [a] | 0.09 | 0.81 | -1.65 | 1.63 |
| Maternal rank | 4.43 | 1.26 | 2.00 | 6.96 |
| **Proportion of infant proximity initiation** | | | | |
| d | 0.07 | 0.14 | -0.22 | 0.33 |
| a | -0.41 | 0.42 | -1.18 | 0.51 |
| b | 3.04 | 0.76 | 1.65 | 4.62 |
| c | 0.23 | 0.28 | -0.34 | 0.74 |

**Table 4 (continued): Response variables**

| Coefficient | Estimate | Standard Error | 95% CI Lower | 95% CI Upper |
|---|---|---|---|---|
| Sex (male) | 2.06 | 1.15 | 0.08 | 4.56 |
| Parity (nulliparous) | 1.32 | 1.52 | -1.43 | 4.58 |
| Fruit availability [a] | 0.27 | 0.92 | -1.63 | 2.07 |
| Maternal rank | 2.89 | 1.29 | 0.51 | 5.58 |
| **Proportion of mother body contact initiation** | | | | |
| d | 1.96 | 0.21 | 1.56 | 2.41 |
| a | 1.03 | 0.25 | 0.57 | 1.55 |
| b | 3.71 | 0.70 | 2.57 | 5.28 |
| c | 0.15 | 0.19 | -0.20 | 0.54 |
| Sex (male) | 2.88 | 1.21 | 0.85 | 5.55 |
| Parity (nulliparous) | 1.51 | 1.44 | -0.99 | 4.63 |
| Fruit availability [a] | -0.01 | 0.88 | -1.89 | 1.62 |
| Maternal rank | 4.61 | 1.30 | 2.25 | 7.31 |
| **Proportion of infant body contact initiation** | | | | |
| d | -0.43 | 0.06 | -0.56 | -0.30 |
| a | 1.19 | 0.56 | 0.13 | 2.36 |
| b | 2.97 | 0.64 | 1.87 | 4.36 |
| c | -0.21 | 0.21 | -0.61 | 0.21 |
| Sex (male) | 2.72 | 1.22 | 0.64 | 5.41 |
| Parity (nulliparous) | 0.91 | 1.55 | -1.83 | 4.18 |
| Fruit availability [a] | -0.05 | 0.86 | -1.87 | 1.58 |
| Maternal rank | 3.20 | 1.47 | 0.40 | 6.15 |

*Note.* [a] Fruit availability was z-transformed with an original mean of 30.291 and a standard deviation of 16.789.

#### Coefficients for the age-trajectory of the transition from dependent to independent feeding and locomotion

**Table 5** Coefficients for the age-trajectory of the transition from dependent to independent feeding and locomotion (mean, standard deviations, and credible intervals of posterior distribution

| Coefficient | Estimate | Standard Error | 95% CI Lower | 95% CI Upper |
|---|---|---|---|---|
| **Proportion of nursing time** | | | | |
| d | -0.14 | 0.21 | -0.52 | 0.30 |
| a | 1.73 | 0.27 | 1.19 | 2.26 |
| b | 2.35 | 0.64 | 1.28 | 3.76 |
| c | -0.10 | 0.20 | -0.47 | 0.30 |
| Sex (male) | 2.87 | 1.14 | 0.98 | 5.42 |
| Parity (nulliparous) | 1.39 | 1.37 | -1.15 | 4.31 |
| Fruit availability [a] | -0.70 | 0.67 | -2.16 | 0.54 |
| Maternal rank | 5.74 | 1.31 | 3.28 | 8.43 |
| **Proportion of independent feeding time** | | | | |
| d | -3.16 | 0.16 | -3.49 | -2.85 |
| a | -0.337 | 0.25 | -0.84 | 0.15 |
| b | 4.65 | 0.72 | 3.43 | 6.26 |
| c | 0.11 | 0.15 | -0.18 | 0.43 |
| Sex | -0.02 | 0.18 | -0.36 | 0.35 |
| Fruit availability [a] | -0.58 | 0.36 | -1.33 | 0.11 |
| **Proportion of carrying time** | | | | |
| d | -1.19 | 0.13 | -1.46 | -0.93 |
| a | 2.57 | 0.16 | 2.25 | 2.88 |
| b | 4.20 | 1.08 | 2.17 | 6.40 |
| c | 0.53 | 0.17 | 0.22 | 0.92 |
| Sex (male) | 3.92 | 1.00 | 2.13 | 6.07 |
| Parity (nulliparous) | 2.23 | 1.37 | -0.29 | 5.13 |

**Table 5 (continued): Response variables**

| Coefficient | Estimate | Standard Error | 95% CI Lower | 95% CI Upper |
|---|---|---|---|---|
| Fruit availability [a] | -0.84 | 1.09 | -2.97 | 1.35 |
| Maternal rank | 5.49 | 1.94 | 1.37 | 8.93 |
| **Proportion of independent locomotion time** | | | | |
| d | -4.23 | 0.44 | -5.22 | -3.50 |
| a | 0.40 | 0.12 | 0.18 | 0.66 |
| b | 2.27 | 0.30 | 1.73 | 2.93 |
| c | 0.36 | 0.13 | 0.11 | 0.64 |
| Sex (male) | -0.00 | 0.09 | -0.17 | 0.17 |
| Fruit availability [a] | 0.14 | 0.05 | 0.05 | 0.24 |

*Note.* [a] Fruit availability was z-transformed with an original mean of 30.270 and a standard deviation of 16.799.
